# Aberrations in stromal signaling, muscle contractility and epithelial architecture underlie poor embryo-implantation outcomes after murine ovarian stimulation

**DOI:** 10.64898/2026.06.14.732145

**Authors:** Harini Raghu Kumar, Manoj Madhavan, Lisa Zou, Curtis Chen, Ryan Yoder, Gregory W. Burns, Emmanuel N. Paul, Nataki C. Douglas, Ripla Arora

## Abstract

Ovarian stimulation is widely used in assisted reproductive technologies, yet its effects on uterine architecture and embryo implantation remain poorly understood. Using a mouse model, we show that ovarian stimulation or superovulation disrupts pre-implantation luminal epithelial folding and induces aberrant smooth muscle structure and contractile function. These structural defects result in embryo trapping within aberrant longitudinal folds, impaired implantation chamber formation, misalignment of the embryo-uterine axis, and subsequent embryo loss. These ovarian stimulation effects were reversible after rest and restoration of normal estrus cycling. Transcriptomic analysis suggests widespread disruption in the stroma and immune compartments of the stimulated uteri. Pathway analysis revealed significant disruption of stromal extracellular matrix and enhanced probability of stroma-immune communication via collagen signaling. Platelet derived growth factor receptor A (PDGFRA) expression was elevated in both the stroma and smooth muscle of the stimulated uteri. Short-term pharmacological inhibition of PDGFRA in the stimulated uteri prior to implantation fully restored epithelial fold transition and implantation chamber formation and partially restored smooth muscle architecture and contractility. Importantly, PDGFRA protein was also elevated in endometrial biopsies from women undergoing ovarian stimulation when compared to natural cycle biopsies. Together, this study establishes muscle contractions and stromal and smooth muscle PDGFRA signaling as novel non-cell autonomous regulators of uterine epithelial architecture critical for embryo implantation.

## INTRODUCTION

Successful embryo implantation occurs in the window of receptivity and is a highly coordinated process between the endometrium and a good-quality embryo. Even a minor alteration in the uterine microenvironment during the receptivity window can disrupt the process of embryo implantation^1^. Defects during early pregnancy can lead to implantation failure and recurrent pregnancy loss and are a major cause of infertility worldwide^2,3^. Assisted reproductive technologies including in vitro fertilization (IVF) are used to treat infertility^4,5^. However, IVF pregnancies are associated with complications including recurrent implantation failure, preterm birth, preeclampsia, and increased perinatal morbidity^6,7^. To improve pregnancy success rates for patients with recurrent implantation failure and after IVF and embryo transfer, it is critical to better understand the endometrial factors that contribute to a successful implantation.

The endometrium undergoes cyclical remodeling as part of the human menstrual cycle consisting of the follicular (proliferative) and luteal (secretory) phases^8^. During the proliferative phase, follicle-stimulating hormone promotes follicle maturation, leading to rising estrogen levels that stimulates endometrial proliferation. In the secretory phase a surge in luteinizing hormone (LH) triggers ovulation, after which progesterone secreted by the corpus luteum prepares the endometrium for implantation. Successful implantation requires precise synchrony between embryo development and uterine maturation, within the window of receptivity which occurs 6–10 days after ovulation^9^. One oocyte is ovulated in a physiologic menstrual cycle. In contrast, ovarian stimulation with exogenous gonadotropins is utilized to induce the simultaneous maturation of multiple ovarian follicles with a goal of harvesting multiple mature oocytes that can be utilized for IVF^10–12^. A robust response to this treatment produces supraphysiological estradiol levels and an early rise in progesterone levels prior to administration of the ovulation trigger^13^. The premature rise in progesterone at 2-5 times physiologic levels^14^ is thought to advance the window of implantation by 1–2 days, leading to premature stromal proliferation and early appearance of progesterone receptor, estrogen receptor, and pinopodes on luminal epithelial surfaces^15,16^. Combined, supraphysiological estrogen and progesterone levels disrupt the peri-implantation uterine environment, leading to dyssynchrony between stromal and glandular maturation^17^ and endometrial transcriptomic changes that affect cellular pathways associated with endometrial receptivity and immune regulation^18–23^. Thus, ovarian stimulation results in a sub-optimal uterine environment that compromises embryo implantation.

Following IVF, high quality embryos can be transferred into the uterus three to five days after the oocyte retrieval as a fresh embryo transfer or they can be vitrified (flash frozen) and cryopreserved for a frozen embryo transfer cycle remote from ovarian stimulation^4^. While embryo quality is comparable between fresh and frozen transfers^24,25^, frozen embryo transfer consistently results in better implantation outcomes, strongly suggesting that the uterine environment could be compromised in fresh embryo transfer cycles^17,24,26–28^. Mouse models have been widely used to investigate the uterine consequences of ovarian stimulation, a condition experimentally induced through superovulation protocols. Similar to human reports of reduced implantation in fresh embryo transfers, mouse studies show greater embryo loss when embryos are transferred into superovulated uteri compared with natural-cycle uteri^29–31^. Further, superovulation mouse models also display supraphysiological hormone levels, aberrant endometrial maturation and impaired uterine receptivity due to abnormal extracellular matrix remodeling and altered immune signaling^32,33^. Architecturally, analysis of thick tissue slices suggests altered vascular remodeling, reduced gland volume and aberrant luminal organization with a deflected luminal axis in superovulation compared to controls^34^. Despite extensive characterization of uterine abnormalities, the direct effect of superovulation on uterine 3D architecture has not been examined.

Uterine contractility during ovarian stimulation in the early luteal phase, when progesterone levels are rising, appears to be more frequent compared to natural cycles^35^. Previous studies in humans have reported that increased contraction activity is negatively correlated with clinical pregnancies^35–37^. Smooth muscle contractility plays an essential role in epithelial sheet remodeling by modulating tissue expansion in organs such as the intestine, lung, and oviduct^38,39^. Koyama et al.^38^ demonstrated that the ratio of epithelial length to surrounding smooth muscle length is a critical mechanical determinant of fold patterning in tubular organs. While supraphysiologic ovarian hormone levels during ovarian stimulation can modulate uterine contractility, how ovarian stimulation affects smooth muscle and how smooth muscle structure and function impacts uterine epithelial remodeling has not been studied.

We recently showed that the uterine epithelium remodels in preparation for embryo implantation^40,41^. At the time of embryo entry into the uterus, the luminal epithelium displays longitudinal folds that run along the uterine oviductal-cervical axis. Prior to embryo dispersal and implantation site formation, the entire epithelium transitions from longitudinal folds to transverse folds that run parallel to the uterine mesometrial-antimesometrial axis. Failure to transition from longitudinal to transverse folds is associated with pregnancy loss at mid-gestation^40–43^. In this study, we demonstrate that superovulation induces a similar failure of preimplantation transition of longitudinal folds to transverse folds, causing aberrant embryo-uterine axis alignment, implantation chamber formation and post-implantation embryo loss. We also observed that superovulation disrupts smooth muscle architecture and uterine contractility during pre-implantation stages. Using transcriptomics and label-free imaging, we show that superovulation heavily impacts the stromal and smooth muscle compartments, resulting in increased PDGFRA expression, extracellular matrix disorganization and excess collagen deposition. Finally, we show that pharmacological inhibition of PDGFRA signaling rescues uterine contractility, preimplantation epithelial folding and implantation chamber formation. Our study highlights novel molecular and structural determinants of uterine receptivity to better understand the embryo-uterine interface during the critical process of embryo implantation.

## RESULTS

### Superovulation disrupts pre-implantation uterine epithelial folding, smooth muscle architecture and uterine contractility

We examined the 3D architecture of the pre-implantation uteri from control and superovulated C57BL/6 mice at gestational day (GD) 3 1200 h (**Fig. 1A**). As previously described, at GD3 1200h control uteri displayed transverse epithelial folds along the M-AM axis (median angle = 32.3°; **Fig. 1B,B’,D**)^40,41^. In contrast, uteri from superovulated mice displayed longitudinal folds along the O-CX axis (median angle = 73.65°; **Fig. 1C,C’,D**). These longitudinal folds are present predominantly near the oviduct and mid-horn, while the cervical region largely displayed transverse folds (**Supplementary Fig. 1A,B,C**). Superovulation produces more embryos than in a natural mating cycle^44,45^. To ensure the excess embryos were not the cause of aberrant uterine folding, we evaluated pseudopregnant superovulated females (generated by mating with vasectomized males). Similar to pregnant superovulated uteri, we observed longitudinal epithelial folds (median angle = 84.6°, **Supplementary Fig. 2B,B’,C**) in the GD3 1200h pseudopregnant superovulated uteri. While uterine folding structural abnormalities were observed, all embryos in superovulated uteri (n = 36) were morphologically at the blastocyst stage, similar to control uteri at GD3 1200h (n = 16) (**Supplementary Fig. 3A**).

**Figure 1.**
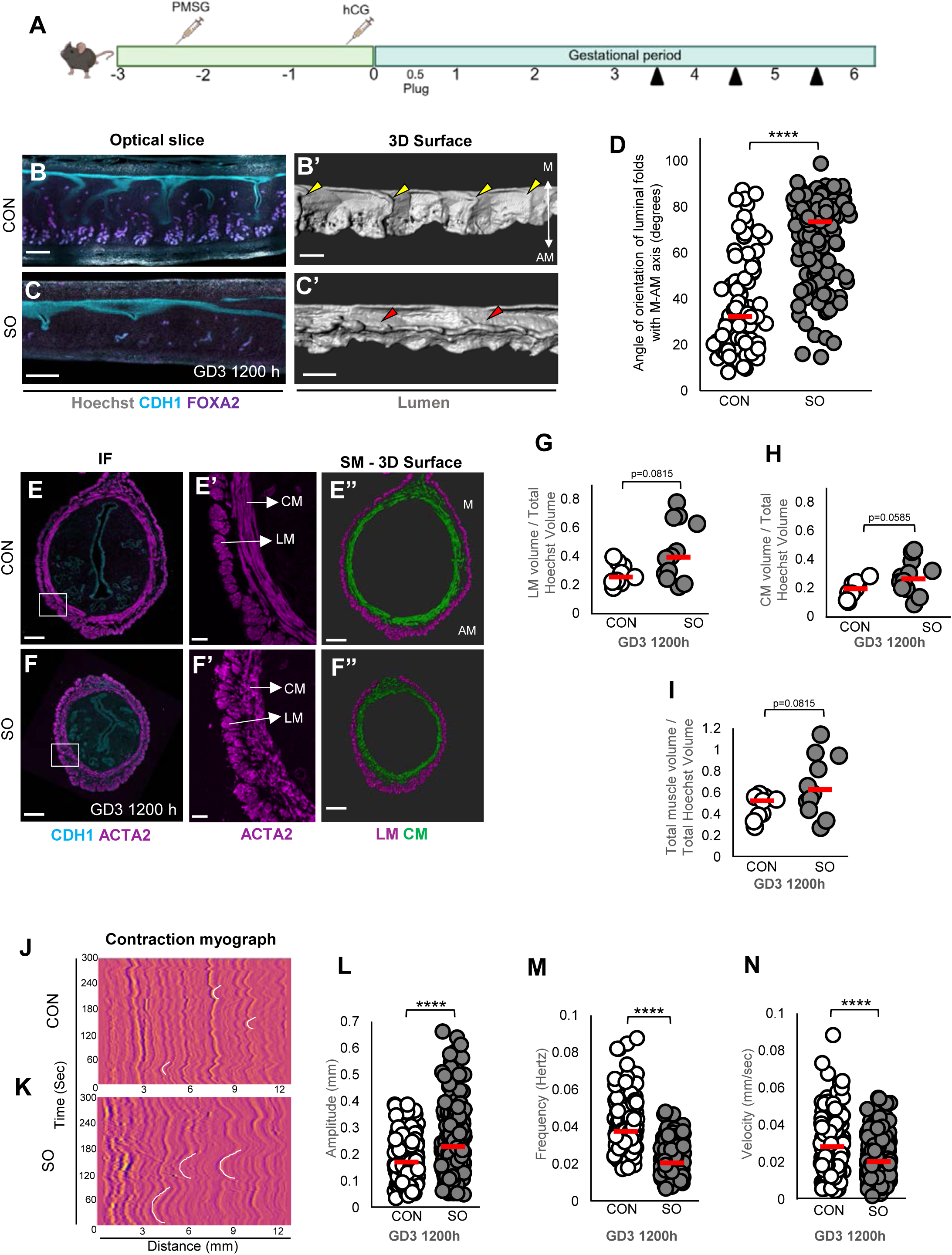
Superovulation leads to failed transition of folds, disrupted smooth muscle architecture, and altered smooth muscle contractions at GD3 1200h. **(A)** Schematic of the superovulation protocol. Black arrowheads: dissection time points. **(B-C)** Optical confocal slices of control **(B)** and superovulated **(C)** uteri. **(B’-C’)** 3D reconstruction of the lumen in **B** and **C** demonstrates transverse folds (yellow arrowheads) in control and longitudinal folds (red arrowheads) in superovulated uteri. **(D)** Quantification of fold angle relative to the M-AM axis in control and superovulated uteri (n = 4 mice/group, *p*<0.05). **(E-F)** ACTA2 immunofluorescence staining in control **(E)** and superovulated **(F)** uteri. **(E’-F’)** Higher magnification of **E** and **F** suggests aberrant smooth muscle in superovulated uteri compared to controls. **(E”-F”)** 3D reconstruction of longitudinal and circular smooth muscle in **E**, **F**. **(G-I)** Quantification of longitudinal **(G)**, circular **(H)**, and total **(I)** smooth muscle volume (n=3 mice/group). Superovulation results in increased volume of the longitudinal and circular muscle compared to controls (*p*< 0.10). **(J-K)** Spatiotemporal plots (myographs) reflecting smooth muscle contractility at in control **(J)** and superovulated **(K)** uteri. **(L-N)** Contraction metrics: **(L)** amplitude, **(M)** frequency, and **(N)** velocity in control and superovulated uteri (*n*=3 mice/group, *p*<0.05). White lines on the myographs indicate example waves. Superovulation induces increased amplitude, decreased frequency and decreased velocity of contractions compared to controls. Statistical test: Mann–Whitney *U*-test. Red dashed line indicates medians. ns, non-significant. M, mesometrial pole; AM, anti-mesometrial pole. CON, control; SO, superovulation; CM, circular smooth muscle; LM, longitudinal smooth muscle. Scale bars: **B,B’,C,C’:** 300 µm; **E,E”,F,F”:** 200 µm; **E’,F**’: 50 µm.

Given that smooth muscle can regulate epithelial organization in tubular organs^39^, we sought to determine if superovulation impacts uterine smooth muscle architecture and function. Immunofluorescence labeling with ACTA2 revealed enlargement and disorganization of both longitudinal and circular uterine smooth muscle fibers in superovulated uteri compared to controls (n=4 mice per group; **Fig. 1E,E’,E”,F,F’,F”).** ACTA2 quantitation revealed a trend towards increased normalized smooth muscle volume of both the longitudinal smooth muscle (Median: control=0.255 normalized unit (nu); superovulated =0.392nu) and circular smooth muscle (Median: control=0.193nu; superovulated =0.262nu) in superovulated uteri at GD3 1200 h (**Fig. 1E”,F”,G–I**), although the data did not reach significance. To determine the function of smooth muscle, we measured uterine contractions using an imaging-based method and generated spatiotemporal plots (myographs) at GD3 1200 h (**Fig. 1J,K, Supplementary Video 1 and 2**)^46^. Superovulated uteri showed a significant increase in contraction amplitude (Median: control=0.1714mm; superovulated =0.2300mm) and a decrease in contraction frequency (Median: control=0.0376Hz; superovulated =0.0206Hz) and velocity (Median: control=0.02842mm/sec; superovulated=0.02033mm/sec) (n = 3 mice/6 horns per group; P < 0.0001, Mann–Whitney U test) (**Fig. 1L-N**). These data suggest that superovulation disrupts luminal epithelial fold transition, smooth muscle structure, and uterine contractility in the pre-implantation uterus.

### Superovulation disrupts implantation chamber formation and embryo-uterine axes alignment

We observed a higher number of embryos^44,45^ in superovulated compared to control uteri at GD4 1800 h (control=8±1.87; superovulated=15.75±2.95) and GD5 1200 h (control=7.75±2.27; superovulated=14.5±6.34) (**Supplementary Table 1**). We also observed an increase in the number of blue dye positive implantation sites at GD4 1800 h in superovulated compared to control uteri (Control: 8 ± 1.87; superovulated: 14.75 ± 3.11, n=4 mice each, p=0.0571; **Fig. 2A,B**). At GD5 1200 h, the number of blue dye positive implantation sites was higher than controls and lower than GD4 1800h superovulated uteri (Control: 7.75 ± 2.27; SO: 11.25 ± 2.86, n=4 mice each) but these differences did not reach statistical significance (**Fig. 2C,D**). Next, we determined the 3D architecture of the implantation sites. At GD4 1800 h, following implantation, 100% of embryos in control uteri were located within flat peri-implantation regions and exhibit normal V-shaped implantation chambers with the embryonic-abembryonic axis aligned along the mesometrial–antimesometrial axis (Ch_norm_; median angle = 12.8°; **Fig. 2E,I**). In contrast, superovulated uteri displayed variability in the implantation chamber formation. 6.25% of the embryos failed to initiate a chamber (Ch_null_; n=4 mice; #embryos = 4/64), and 39.06% of embryos in the superovulated uteri were trapped within longitudinal folds (Ch_trap_; n=4 mice; # embryos = 25/64; **Fig. 2F,G**). These trapped embryos formed abnormal implantation chambers and their embryonic-abembryonic axis was oriented parallel to the maternal oviductal-cervical axis (median angle = 86.3°; P<0.001; **Fig. 2I**) (**Supplementary Fig. 3B, Panels 2–3**^41^). The remaining 54.68% of embryos in superovulated uteri formed normal V-shaped implantation chambers (Ch_norm_; n=4 mice; # embryos 35/64), displayed proper embryo–uterine alignment (median angle = 19.7°; **Fig. 2H,I**), and were comparable in size and morphology to control embryos (**Supplementary Fig. 3B, Panel 4**). Importantly, embryos trapped in longitudinal folds showed blue dye positive reaction similar to those that initiated a Ⅴ-shaped chamber^41,42^.

**Figure 2.**
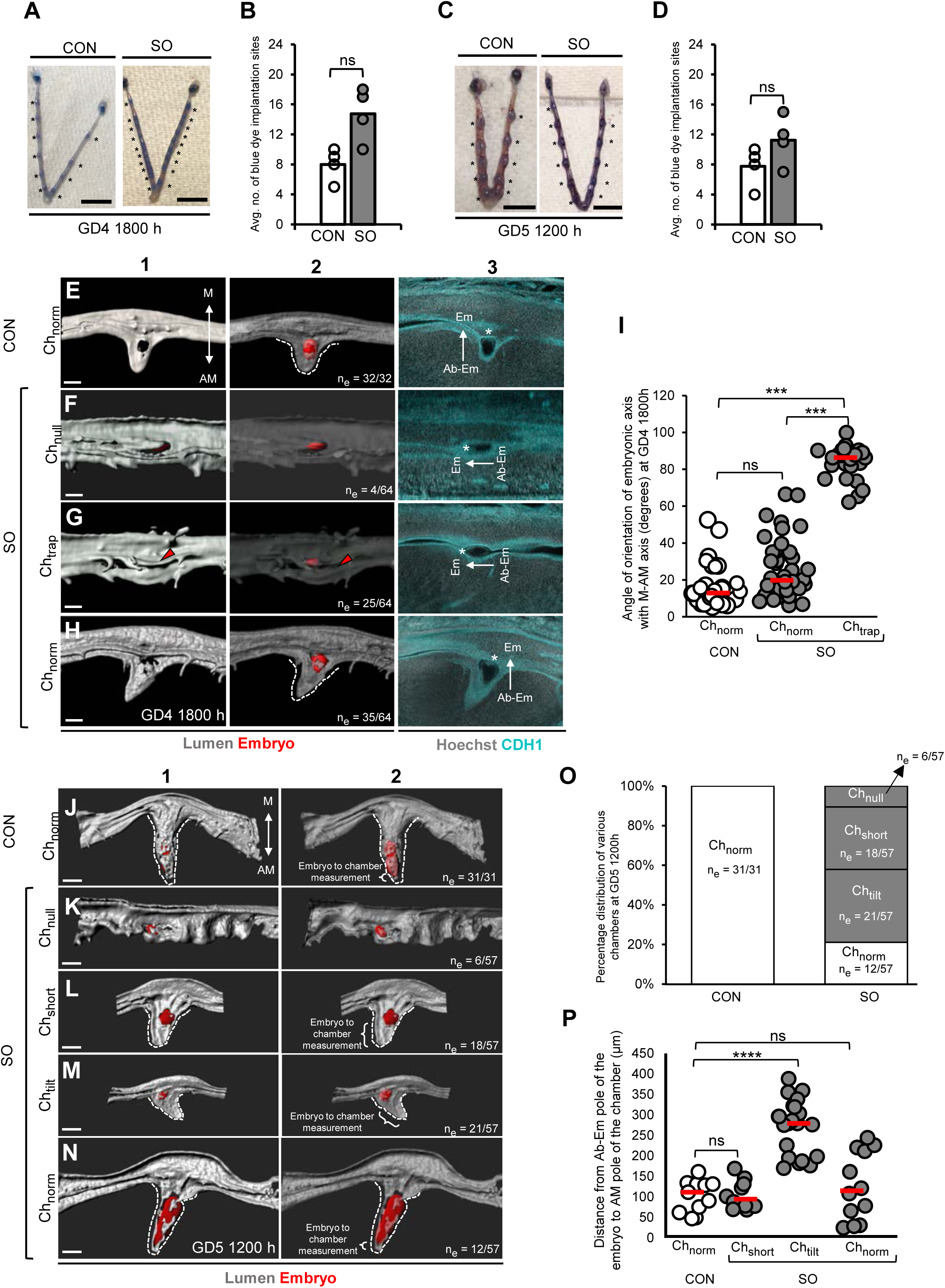
Superovulation disrupts implantation chamber formation and post-implantation embryo-uterine axes alignment. **(A-D)** Implantation site analysis using blue dye injection at GD4 1800h **(A, B)** and GD5 1200h **(C, D)**. Asterisk: blue dye positive implantation sites. **(E-H)** 3D surface and optical slice views of implantation chambers at GD4 1800 h in control **(E)** and superovulated **(F-H)** uteri. Control uteri show a normal V-shaped chamber with an aligned embryo (Ch_norm_). Superovulated uteri show variations in epithelial morphology at the embryo-lumen interface: **(F)** failure to initiate a chamber (Ch_null_), **(G)** embryo trapped in longitudinal fold (Ch_trap_) and **(H)** normal V-shaped chamber (Ch_norm_). Panel 1: 3D lumen surface (gray). Panel 2: transparent 3D lumen and embryo surface (red). Panel 3: Frontal optical slice view of the embryos. Asterisks: inner cell mass; red arrowhead: longitudinal fold. **(I)** Quantification of embryo orientation relative to the M–AM axis in control (n = 4 mice, n_e_ = 32 embryos) and superovulated (n = 4 mice, n_e_ = 64 embryos) uteri at GD4 1800 h. *p* < 0.05, Kruskal-Wallis test with Dunn’s multiple comparison. **(J-N)** 3D surface view of implantation sites at GD5 1200 h in control **(J)** and superovulated **(K-N)** uteri. Control uteri show an elongated V-shaped chamber. Superovulated uteri display variability in chamber formation: **(K)** no chamber (Ch_null_), **(L)** short chamber (Ch_short_), **(M)** tilted chamber (Ch_tilt_), and **(N)** normal elongated chamber (Ch_norm_). Panel 1: 3D lumen surface (gray). Panel 2: transparent 3D lumen and embryo (red) surface. **(O)** Percentage distribution of chamber types in control and superovulated uteri at GD5 1200 h. **(P)** Quantification of distance from abembryonic pole of the embryo to the anti-mesometrial pole of the chamber in control (n = 4 mice, n_e_ = 31 embryos) and superovulated (n = 4 mice, n_e_ = 57 embryos) uteri at GD5 1200 h. *p* < 0.05, Welch’s t-test. Red dashed lines on graphs indicate the median. ns, non-significant; M, mesometrial pole; AM, anti-mesometrial pole; Em, embryonic pole; Ab-Em, abembryonic pole. n_e,_ Number of embryos; CON, control; SO, superovulation. Scale bars: **A, C:** 5mm; **E,F,G,H:** 100 μm; **J,K,L,M,N:** 200 μm.

At GD5 1200 h, 100% of embryos in control uteri formed elongated V-shaped implantation chambers (Ch_norm_, **Fig. 2J,O**). In contrast, embryos in superovulated uteri showed variability in implantation chamber formation, which we categorized into four distinct groups. 10.5% of embryos were visibly smaller (**Supplementary Fig. 3C, Panel 2**) and failed to initiate an implantation chamber (Ch_null_; n=4 mice; # embryos = 6/57) **(Fig. 2K,O**). 31.5% of embryos formed short, straight chambers (**Fig. 2L,O**), with embryos at an early egg cylinder stage (Ch_short_; n=4 mice; # embryos = 18/57). 36.8% of embryos developed intermediate-sized chambers with a tilt (**Fig. 2M,O**) (Ch_tilt_; n=4 mice; # embryos = 21/57). Ch_tilt_ embryos appeared small and had a misaligned embryo-uterine axis within the chamber (**Supplementary Fig. 3C, Panel 3**). Finally, 21% of embryos formed elongated straight chambers (Ch_norm_; n=4 mice; # embryos = 12/57) similar to those observed in control uteri (**Fig. 2N,O**) and embryo morphology resembled normal egg cylinder-stage embryos (**Supplementary Fig. 3C, Panel 4**). Of note, while Ch_norm_ and Ch_tilt_ chambers correlate with blue dye positive implantation sites, Ch_null_ and Ch_short_ chambers failed to show a blue dye positive implantation site (**Fig. 2A-D**). Further qualitative assessment suggested that embryos located within Ch_short_ and Ch_norm_ displayed symmetric decidual-epiblast spacing (**Supplementary Fig. 4B**)^41,47^. In contrast, embryos within Ch_tilt_ exhibited asymmetric spacing between the decidua and epiblast of the egg cylinder embryo, with embryos displaced towards the M-pole, resembling phenotypes observed in other mouse models with defective uterine folding (**Supplementary Fig. 4B**^41^). Thus, while a subset of embryos in superovulated uteri initiated normal chamber formation, nearly 80% embryos displayed implantation chamber defects, ranging from incomplete or absent chamber formation to abnormal chamber orientation and embryo-uterine misalignment.

### Superovulated uteri display post-implantation embryo loss

To determine the fate of embryos exhibiting abnormal implantation-chamber morphology, we examined pregnancies at mid-gestation (GD13 1200 h). The average number of embryos (live embryos and resorption sites combined) was comparable between control mice (8.6 ± 3.14, n = 5 mice; **Supplementary Fig. 3D and Supplementary Table 1**) and superovulated mice (7.4 ± 1.74, n = 5 mice; **Supplementary Fig. 3E and Supplementary Table 1**). However, the mid-gestation (GD13 1200h) embryo count in superovulated uteri was substantially lower (34.37%) than the embryo numbers observed at both GD4 1800 h (superovulation: 16 ± 2.95) and GD5 1200 h (superovulation: 14.25 ± 6.34), indicating that substantial embryo loss occurred between GD4 1800h and GD13 1200 h (**Supplementary Table 1**). These losses did not manifest as resorption sites suggesting that many embryos failed early in post-implantation development.

### Resumption of normal estrus cycling restores uterine architecture in previously superovulated mice

To evaluate whether the uterine structural abnormalities induced by superovulation are reversible over time, we examined pregnancy outcomes in mice that were rested to allow recovery from the effects of ovarian stimulation. Superovulated females were mated to vasectomized males to induce pseudopregnancy and then rested for a 4-week recovery period (∼5-6 estrus cycles) before being set up for natural mating. Uteri from these rested females displayed normal transverse folding at GD3 1200h (n=4 mice, **Supplementary Fig. 5A,A’, B,B’**), proper implantation chamber formation and embryo-uterine alignment at GD4 1800 h (n=4 mice; **Supplementary Fig. 5C,D**), and 100% of embryos were viable at GD13 1200h (n=7 mice; **Supplementary Fig. 5E,F**). Together, these findings show that the detrimental effects of superovulation on uterine structure and implantation are temporary and fully reversible after normal-estrus cycling resumes.

### Single-cell sequencing reveals alterations in epithelial and non-epithelial compartments of the pre-implantation superovulated uterus

To investigate the molecular basis of the structural defects observed during superovulation, we performed single-cell RNA sequencing (scRNA-seq) on whole reproductive tracts collected from control and superovulated mice. Since scRNA-seq captures mRNA differences, we performed the tissue collection at GD3 0900 h (n = 2 mice per group), a time point where in control mice the transition of longitudinal to transverse folds has initiated but is not yet complete^41^. A total of 39,439 cells passed quality control, with an average sequencing saturation of 65.5%, corresponding to approximately 35,000 reads per cell. Uniform Manifold Approximation and Projection (UMAP) analysis of scRNA-seq revealed 22 major cell clusters across both control and superovulated uteri (**Fig. 3A,B**). Cluster identities were assigned based on the expression profiles of canonical cell-type markers (**Fig. 3C**). Compared to controls the relative proportion of dendritic cells and T cells increased in the superovulated uteri by 35.7% and 40.21% respectively (**Fig. 3D**), while all other cell-types were comparable between the two conditions. When evaluating the stromal compartment, we identified two distinct sub-clusters – deep stroma and superficial stroma^48^. Deep stroma uniquely expressed *C3*, *Postn* and *Clec3b*, and superficial stroma uniquely expressed *Hoxa11*, *Sfrp2*, *Hoxa10*, *Fst*, *Hsd11b2*, and *Cdh11* (**Fig. 3C**). Expression of higher levels of *Mki67*^48^, *Hand2*^48^, *Pgr*^48^ and *Hand2os1*^49^ suggests that the superficial stromal cluster is the sub-epithelial stroma (**Fig. 3E’,F’,G,H**), whereas the deep stroma compartment is farther away from the epithelium and closer to the myometrium (**Fig. 3E,F**).

**Figure 3.**
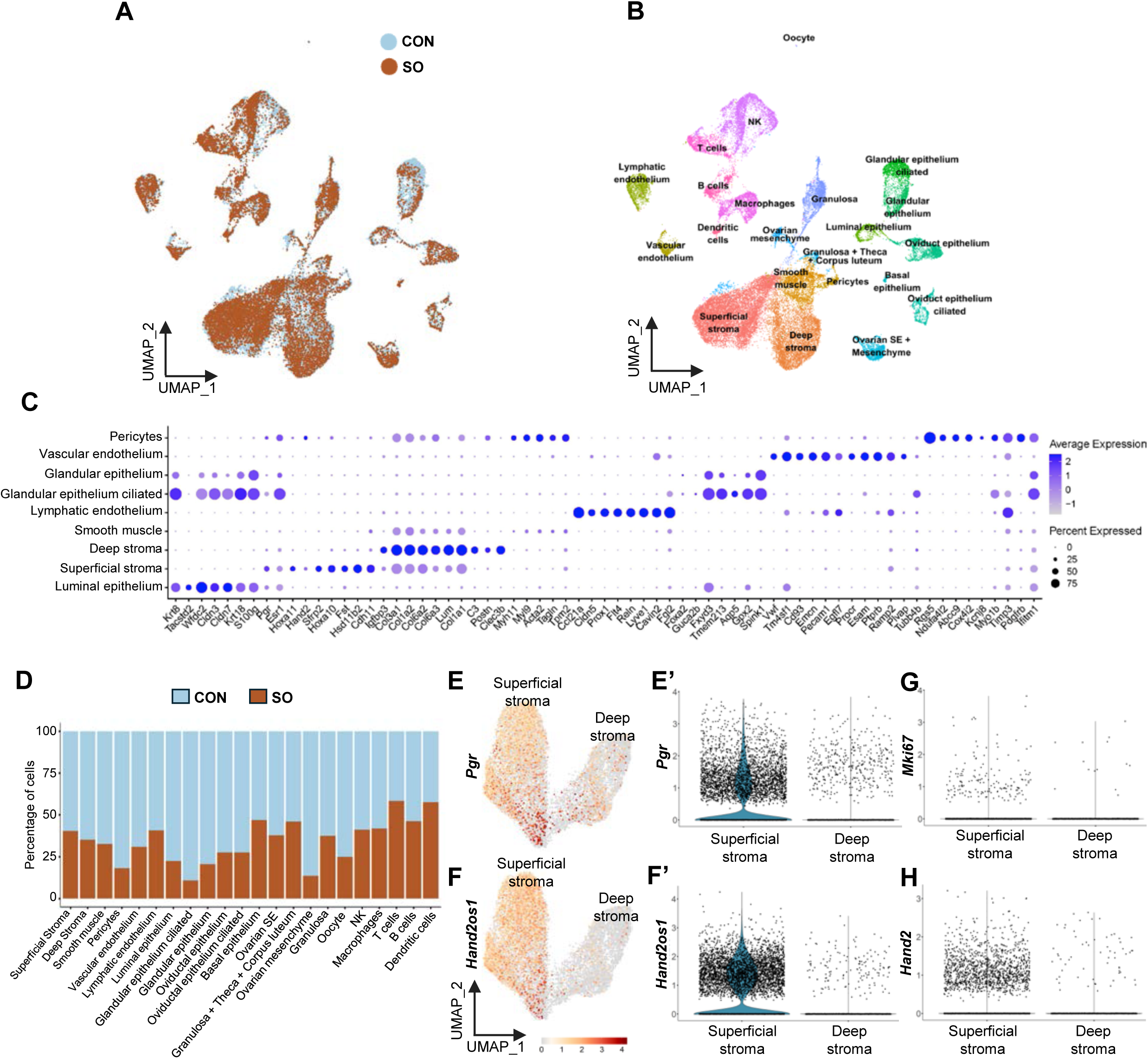
Pre-implantation transcriptomic profiling and cell population characterization by single-cell RNA sequencing. **(A)** Uniform Manifold Approximation and Projection (UMAP) of isolated cells in the mouse uterus by single-cell RNA sequencing at GD3 0900 h (n = 2 mice/group), showing integration of control and superovulated uterine cells with no gain or loss of clusters due to superovulation. **(B)** UMAP visualization of cell clusters identified. **(C)** Dot plot showing the expression of cell type-specific marker genes used for cluster identification. **(D)** Stacked bar plot showing the proportion of cells within each cluster. Superovulated uteri show an increase in dendritic cells and T cells. **(E-F)** UMAP showing *Pgr* **(E)** and *Hand2os1* **(F)** expression, identifying the two stromal sub-clusters: superficial stroma (sub-epithelial stroma), and deep stroma (adjacent to the smooth muscle). **(E’-F’,G-H)** Violin plots showing high *Pgr* **(E’)**, *Hand2os1* **(F’)**, *Mki67* **(G)**, and *Hand2* **(H)** expression in the superficial stroma compared to the deep stroma. CON, control; SO, superovulation.

Since the luminal epithelium displayed a defect in folding, we first evaluated differentially expressed genes (DEGs) in this compartment. We identified 62 DEGs in the luminal epithelium and of these, 8 genes implicated in cell autonomous epithelial folding, including *Actb*, *Actg1*^50^, *Myl12a*^51^, *Pfn1*^52^, *Dstn*^53^, *Rdx*, and *Cnn3*^54^ were downregulated **(Fig. 4A** and **Supplementary Fig. 6**). However, these genes were not unique to the epithelial compartment and were widely expressed in multiple uterine cell types (**Supplementary Fig. 6**). We further evaluated other compartment DEGs to determine superovulation-induced gene signatures. Amongst the immune cells, while there was an increase in the number of dendritic and T-cells in the superovulation condition (**Fig. 3D**), we observed no DEGs in the dendritic cells and only 30 DEGs in the T-cells. The Natural killer (NK) cell gene signature was most dysregulated due to superovulation with 1047 DEGs (**Fig. 4B**). Intriguingly, we observed 467 DEGs in the deep stroma and 1005 DEGs in the superficial stroma (**Fig. 4C,D**). We also observed 79 DEGs in the smooth muscle layer (**Fig. 4E**). Notably, Platelet-derived growth factor receptor alpha (*Pdgfra*), a stromal marker known to regulate epithelial folding in the intestine^55^, was upregulated in the deep stroma, the superficial stroma and the smooth muscle compartment (**Fig. 4C-E and Supplementary Fig. 7A-C**).

**Figure 4.**
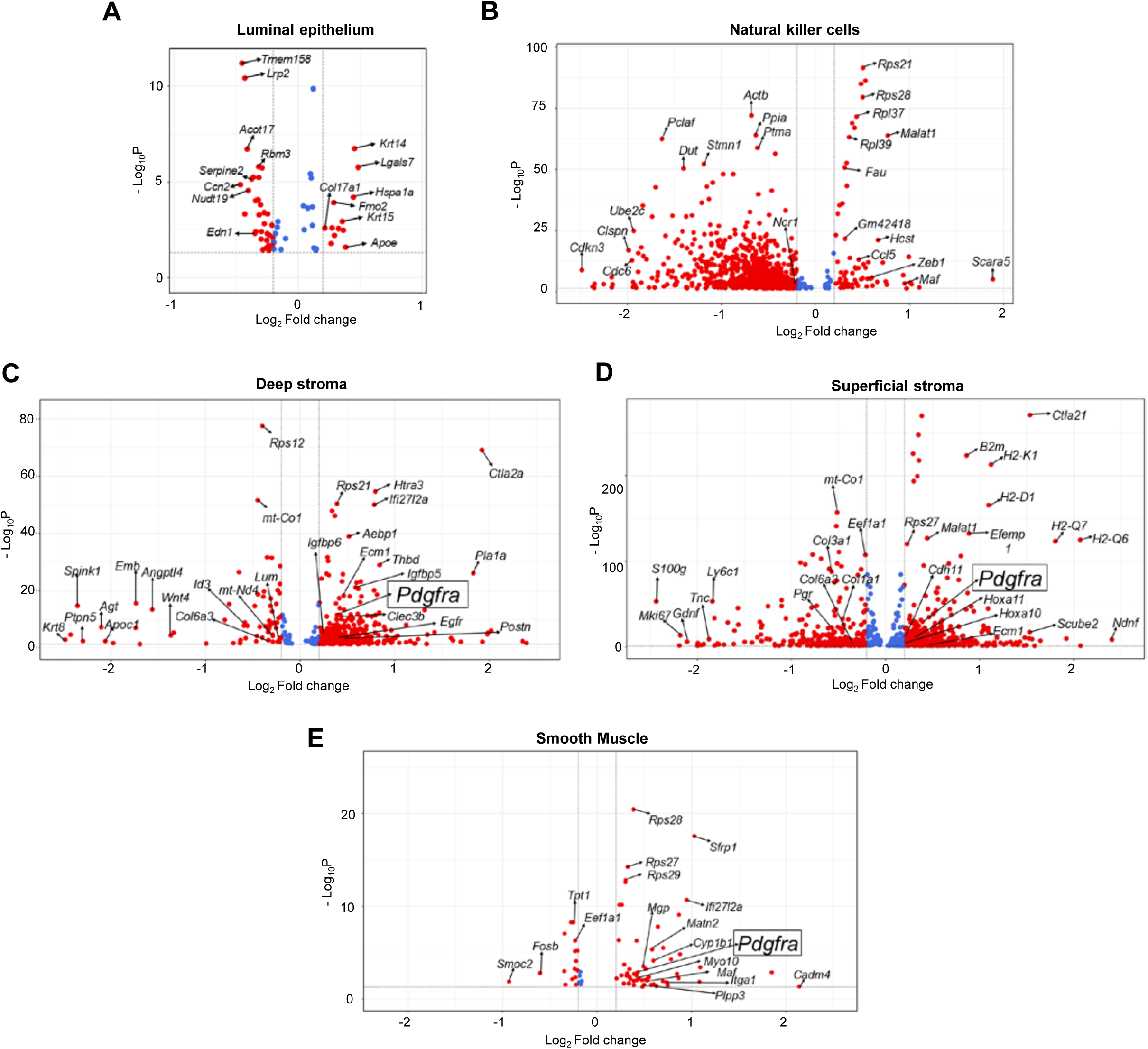
Superovulation induces dysregulation in the pre-implantation luminal, stromal, smooth muscle, and immune compartments. **(A-E)** Volcano plots of differentially expressed genes (DEGs) comparing control and superovulated uteri in the luminal epithelium **(A)**, Natural killer cells **(B)**, deep stroma **(C)**, superficial stroma **(D)** and smooth muscle **(E)**. Red dots: significant DEGs; blue dots: non-significant genes. *Platelet-derived growth factor receptor alpha (Pdgfra)*, highlighted in the black box, is significantly upregulated in both stromal compartments **(C, D)** and smooth muscle **(E)**. CON, control; SO, superovulation.

### Disrupted extracellular matrix organization and increased collagen deposition in pre-implantation superovulated uteri

To further characterize the molecular pathways that differ between control and superovulated stroma, we examined gene ontology (GO) biological process terms of DEGs. Enriched terms in superficial stroma included epithelial cell proliferation and gland development, whereas enriched terms in deep stroma included muscle cell differentiation and striated muscle cell differentiation, further supporting the proximity of the superficial stroma to the epithelial compartment and the proximity of the deep stroma to the uterine smooth muscle. GO biological process terms in both the superficial and deep stroma revealed significant enrichment of extracellular matrix (ECM)-related pathways, including ECM organization, external encapsulating structure organization, and extracellular structure organization (**Fig. 5A,B**). Consistent with this, ECM gene *Ecm1* was upregulated in both stromal compartments and *Efemp1* was upregulated specifically in the superficial stromal compartment (**Fig. 4C,D**). To validate ECM changes reflected by scRNA-seq analysis and to determine collagen organization, we used second harmonic generation (SHG) imaging to detect fibrillar collagen. SHG imaging confirmed enhanced collagen deposition in both the stroma and the smooth muscle of superovulated uteri compared to controls (**Fig. 5C, D**).

**Figure 5.**
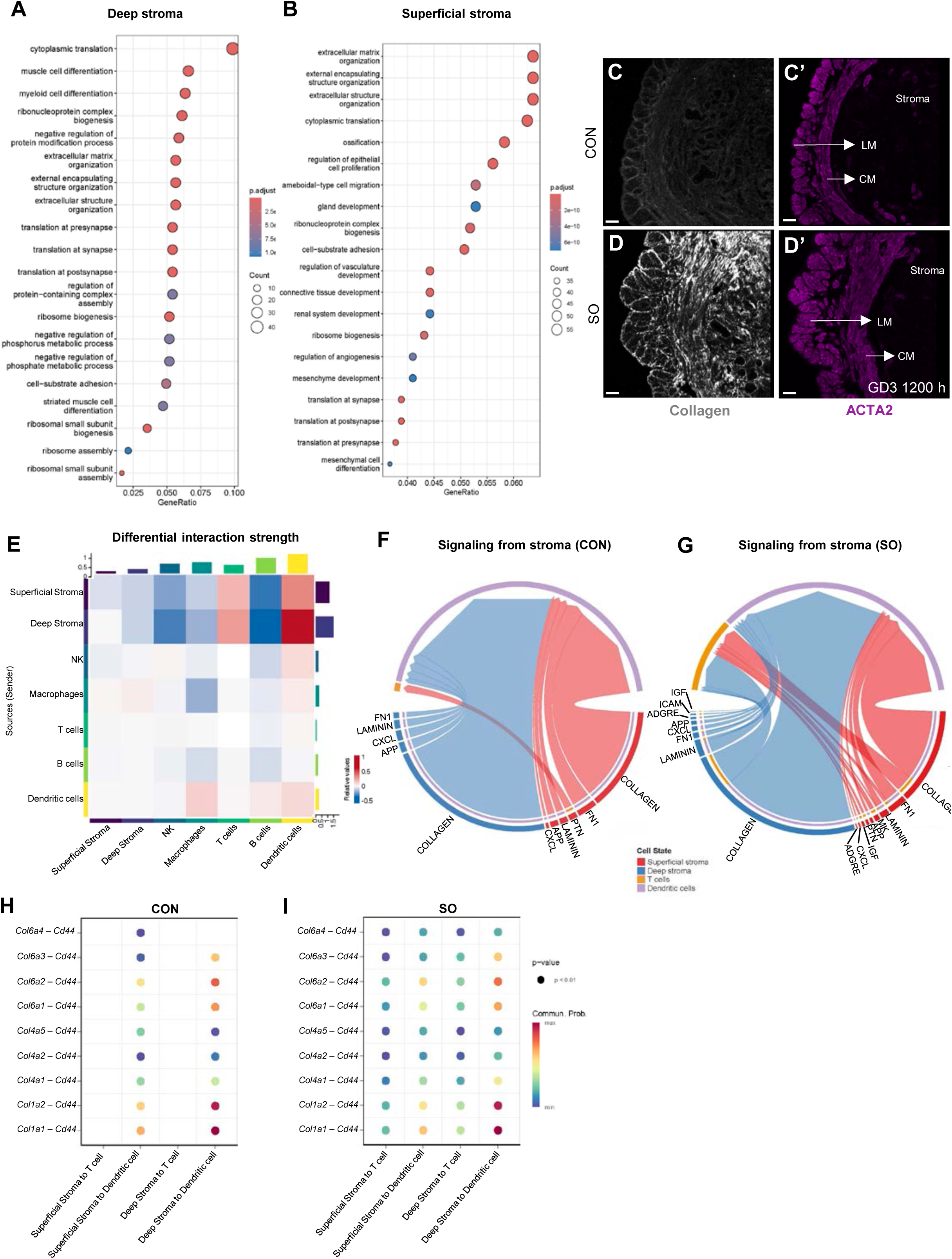
Superovulation induces extracellular matrix dysregulation, enhanced collagen deposition, and amplified stromal-immune cell communication in early pregnancy. **(A-B)** Dot plots depicting the top 20 enriched Gene Ontology Biological Processes terms in the deep stroma **(A)** and superficial stroma **(B)** of superovulation vs control condition. Dot size represents the number of genes associated with each term, and color intensity indicates the adjusted *p*-value. Extracellular matrix organization-related terms are prominently enriched in both stromal compartments after superovulation. **(C-D)** Second harmonic generation imaging of control **(C)** and superovulated **(D)** uteri at GD3 1200 h reveals enhanced fibrillar collagen in the stroma and smooth muscle of superovulated uteri. **(C’-D’)** ACTA2 immunofluorescence staining corresponding to **C** and **D**. **(E)** CellChat analysis showing ligand-receptor-based intercellular signaling between the stroma and immune compartments at GD3 0900 h in control vs superovulation condition. Colors represent the relative signaling strength of each pathway across clusters, with blue indicating decreased communication and red indicating increased communication. **(F-G)** Chord diagrams depicting ligand-receptor-based signaling pathways in control **(F)** and superovulated **(G)** uteri suggests expanded COLLAGEN signaling between stromal cells and T cells. Arc size represents the relative signaling strength of each pathway. **(H-I)** Bubble plots depicting collagen ligand-receptor interactions (*Col–Cd44*) between the superficial stroma, deep stroma, dendritic cells, and T cells in control **(H)** and superovulated **(I)** uteri. CON, control; SO, superovulation; CM, circular smooth muscle; LM, longitudinal smooth muscle. Scale bars: **C,C’,D,D’:** 50 µm.

Using CellChat analysis, we predicted heightened ligand-receptor-based intercellular signaling between the stromal compartment and both the dendritic cells and the T-cells of superovulated uteri (**Fig. 5E**). In addition to increased collagen deposition and disrupted ECM signatures in the superovulated stroma (**Fig. 5A - D**), CellChat revealed an increase in COLLAGEN signaling between deep and superficial stroma and dendritic cells and newly predicted COLLAGEN signaling between deep stroma and T cells in superovulated condition (**Fig. 5G,H,I and Supplementary Fig. 8**). Together, these findings reveal that superovulation induces broad transcriptomic dysregulation across multiple uterine compartments, characterized by ECM remodeling in the stromal and smooth muscle compartments and amplified collagen based stromal-immune communication, pointing to a disrupted uterine microenvironment during the pre-implantation period.

### Superovulation causes elevated PDGFRA protein expression in mouse and human endometrium

scRNA-seq DEG analysis revealed significant upregulation of *Pdgfra* mRNA in superovulated uteri (**Fig. 4C-E**) and stromal PDGFRA is known to modulate epithelial folding in the mouse intestine^55^. An increase in *Pdgfra* mRNA could either indicate that there were more stromal cells expressing *Pdgfra* or that individual cells were expressing higher levels of *Pdgfra*. Further analysis of the scRNA-seq suggests that the level of *Pdgfra* expression per cell is higher in the superovulated superficial and deep stroma (**Supplementary Fig. 7**). To confirm this observation at the protein level, we performed PDGFRA immunofluorescence and determined two metrics: a) the normalized volume of PDGFRA expressing cells and b) the normalized volume of cells expressing high and low levels of PDGFRA based on fluorescence intensity. We first evaluated PDGFRA expression at GD1 1200 h, where both control and superovulated uteri exhibited longitudinal folds (**Fig. 6A,B**). At this stage, neither the number of cells expressing PDGFRA nor the intensity of PDGFRA was significantly different between control and superovulated uteri (**Supplementary Table 2**, **Fig. 6C, D, E, F**, n=3 mice/group). This was true for both the stromal compartment and the smooth muscle compartment. At GD3 1200 h, when control uteri typically transition from longitudinal to transverse folds, there was no difference in overall volume of stromal or smooth muscle cells expressing PDGFRA (**Fig. 6G,H; Supplementary Table 2**). However, the volume of cells expressing PDGFRA at a higher intensity was increased in both stroma and smooth muscle from the superovulated uteri when compared to control (n=3 mice per group; **Fig. 6I,J,K,L**). Additionally, there was a reduction in the volume of cells expressing lower levels of PDGFRA in the stromal compartment. Thus, superovulated pre-implantation uteri display an expansion of high-intensity PDGFRA-expressing cells in the stroma and smooth muscle.

**Figure 6.**
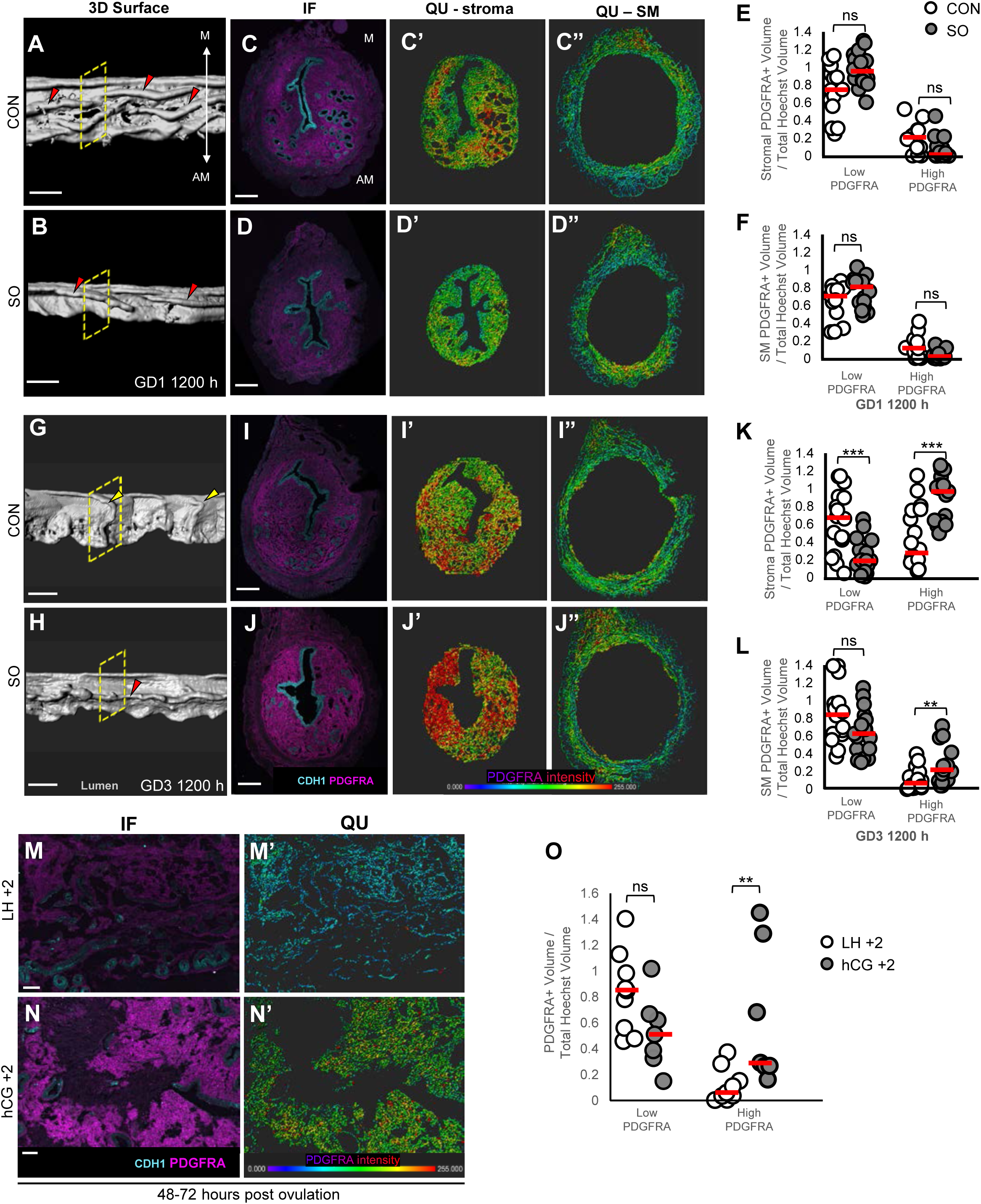
Ovarian stimulation induces elevated PDGFRA expression in the mouse and human uterus. **(A,B)** 3D reconstruction of the lumen and **(C-D)** transverse sections in control **(A,C)** and superovulated **(B,D)** uteri at GD1 1200 h. **(C’-C”, D’-D”)** PDGFRA intensity mapping of the stroma and smooth muscle at GD1 1200 h in control **(C’–C”)** and superovulated **(D’–D”)** uteri (n = 3 mice/group, n_t_ = 14-18 transverse sections). **(E-F)** PDGFRA intensity quantification in stroma **(E)** and smooth muscle **(F)** corresponding to **C’,C”**,**D’,D”**. **(G,H)** 3D reconstructions of the lumen and **(I-J)** transverse sections in control **(G,I)** and superovulated **(H,J)** uteri at GD3 1200 h. **(I’-I”, J’-J”)** PDGFRA intensity mapping of the stroma and smooth muscle in control **(I’–I“**) and superovulated **(J’–J”)** uteri (n = 3 mice/group, n_t_ = 17-18 transverse sections). **(K-L)** PDGFRA intensity quantification in stroma **(K)** and smooth muscle **(L)** corresponding to **I’,I”,J’,J”**. **(M-N)** PDGFRA immunofluorescence staining on human uterus biopsy samples from normal cycling women at LH+2 **(M)** and stimulated women at hCG+2 **(N)**. **(M’-N’)** PDGFRA intensity mapping of stroma in **M** and **N** demonstrates elevated PDGFRA levels in stimulated women (n = 3 samples/group; n_t_ = 7-10 transverse sections). **(O)** PDGFRA intensity quantification of stroma corresponding to **M** and **N**. Statistical test: *p* < 0.05, Mann–Whitney U-test; ns, non-significant; n_t_, number of transverse sections; IF, immunofluorescence; QU, PDGFRA intensity map. PDGFRA signal intensity ranges: in mouse - low <200, high >200; in human - low <128, high >128. QU color scale: purple, low intensity; red, high intensity. Red dashed lines indicate the median. CON, control; SO, superovulation. Scale bars: **A, B, G, H -** 3D surface: 300 µm; **C-C”,D-D”,I-I”,J-J”** - IF and QU: 200 µm, **M,M’,N,N’:** 100 µm.

To determine if endometrial PDGFRA expression is altered by ovarian stimulation in women, we evaluated bulk RNA sequencing data from stage-matched endometrial samples obtained in the peri-ovulatory period and mid-secretory phase of unmedicated, ovulatory menstrual cycles (based on endogenous LH surges) and ovarian stimulation cycles without fresh embryo transfer (based on timing of the human chorionic gonadotropin (hCG) ovulation trigger). For the peri-ovulatory samples, endometrial biopsies were performed 2 days after an LH surge (LH+2) or human hCG trigger (hCG+2) and for the mid-secretory phase samples, endometrial biopsies were performed 9 days after the LH surge (LH+9) or hCG trigger (hCG+9)^17^. We observed that *PDGFRA* mRNA was significantly upregulated in hCG+2 endometrium as compared to LH+2, but *PDGFRA* mRNA levels were comparable in LH+9 and hCG+9 endometrium (**Supplementary File 1 and 2**). In agreement with this, PDGFRA protein expression was significantly elevated during the peri-ovulatory period, in the hCG+2 group compared to the LH+2 group (p<0.01, n = 3 per group; **Fig. 6M,N,O**). We did not observe differences in PDGFRA protein expression in endometrium from hCG+9 compared to LH+9 that coincides with the mid-secretory phase (**Supplementary Fig. 9A,B**). Thus, PDGFRA levels were higher in ovarian stimulation conditions in both mice and humans 48-72 hours after the LH surge, correlating with rising progesterone levels and not the window of receptivity (**Supplementary Table 3**^17^).

### Pharmacological inhibition of PDGFRA signaling rescues pre-implantation folding, muscle contractility and implantation chamber formation in superovulated mice

To determine the functional relevance of increased PDGFRA after superovulation, we examined whether inhibiting PDGFRA activity can rescue the aberrant phenotypes observed in the peri-implantation superovulated uteri. We used imatinib, which is known to inhibit PDGFRA, PDGFRB, and KIT^56^. Interrogation of our scRNA-seq dataset revealed that while *Kit* was minimally expressed in the stroma and smooth muscle, *Pdgfrb* was expressed in the stromal compartment (**Supplementary Fig. 10**). Thus, we used an additional inhibitor of PDGFRA signaling, avapritinib, which inhibits only PDGFRA and KIT^57^. Superovulated mice received an intraperitoneal injection of either imatinib, avapritinib or their respective vehicles at GD2 2200 h (**Fig. 7A**) and the tissue architecture was evaluated at GD3 1200 h to assess folding. Superovulated mice treated with vehicles displayed longitudinal folds at GD3 1200 h (DI-water vehicle, median angle = 64.5°; DMSO+saline vehicle, median angle = 63°; **Fig. 7B,B’,D,D’,F**; **Fig. 1C’**, *p* < 0.05). Strikingly, superovulated mice treated with either imatinib (n=6 mice; median angle = 35.6°; **Fig. 7C,C’,F**) or avapritinib (n=5 mice; median angle = 42.35°; **Fig. 7E,E’,F**) exhibited transverse folds, indicating a complete rescue of pre-implantation folding (**Supplementary Fig. 11**). Importantly, embryos from imatinib treated superovulated mice were at the blastocyst stage and appeared healthy (inset in **Fig. 7C**) suggesting that the treatment did not compromise embryo quality. A subset of embryos in avapritinib-treated superovulated uteri appeared morphologically abnormal (inset in **Fig. 7D,E**). Given the adverse effect of avapritinib on embryo quality, we performed subsequent rescue analysis using Imatinib.

**Figure 7.**
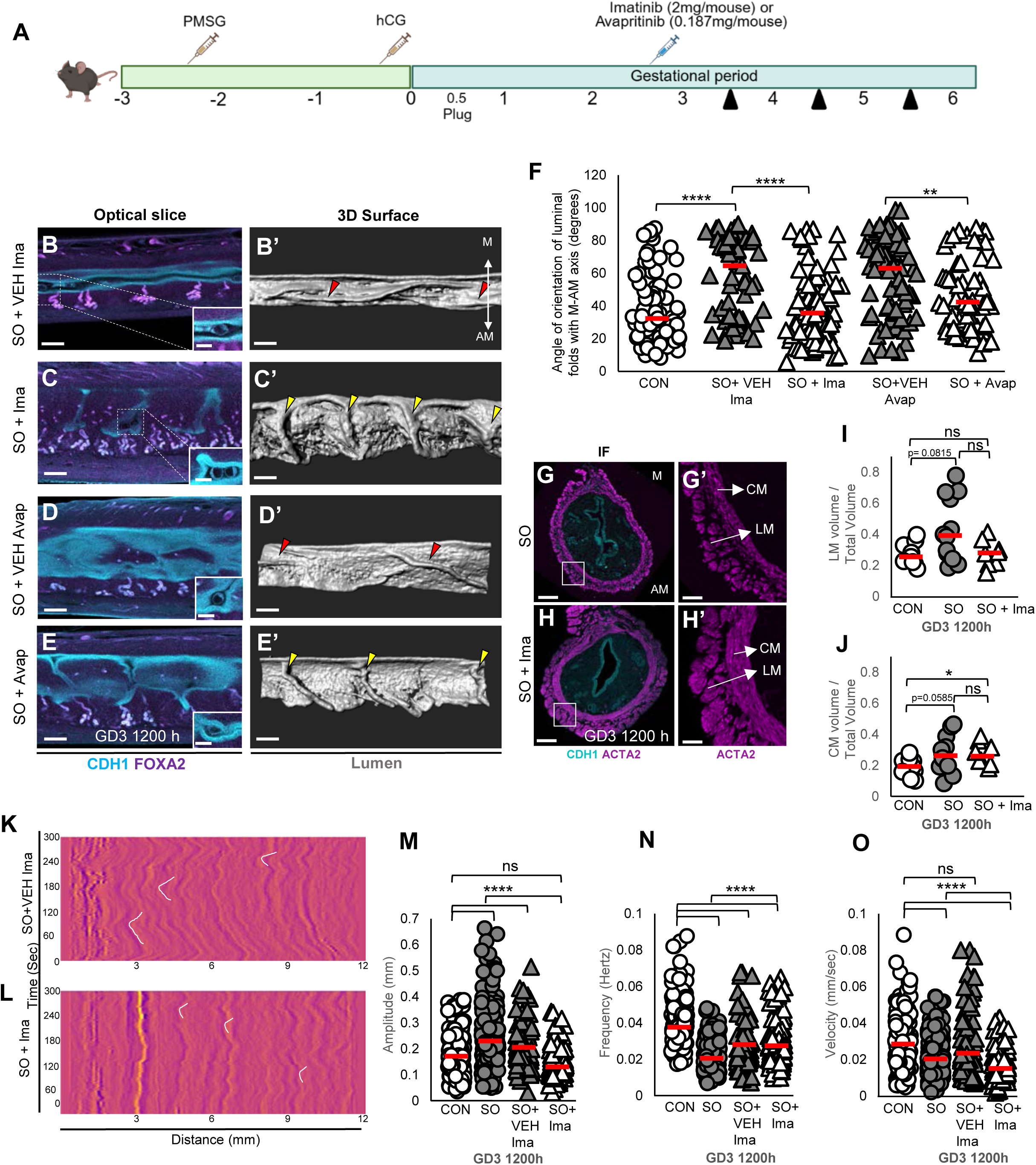
Superovulated mice treated with PDGFRA inhibitor display complete rescue of pre-implantation transverse epithelial folding and partial rescue of smooth muscle structure and contractility. **(A)** Schematic of the superovulation with inhibitor treatment protocol. Black arrowheads: dissection time points. **(B-E)** Optical confocal slices of superovulated mice treated with vehicle or inhibitor at pre-implantation: **(B,C)** superovulation + vehicle/imatinib, **(D,E)** superovulation + vehicle/avapritinib. The inset shows a magnified view of a blastocyst in **B**-**E. (B’-E’)** 3D reconstructions of the lumen shown in **B**, **C, D** and **E.** Yellow arrowheads: transverse folds in superovulated mice treated with inhibitor; red arrowheads: longitudinal folds in superovulated mice treated with vehicles. **(F)** Quantification of fold angle relative to the M–AM axis in control mice, superovulated mice treated with vehicle or inhibitor at GD3 1200 h (*p*<0.05; control, superovulation + imatinib/vehicle: n = 3 mice, superovulation + avapritinib/vehicle: n=5 mice/group). **(G,H)** ACTA2 immunofluorescence staining in superovulated uteri **(G)** and superovulation+imatinib uteri **(H)** at GD3 1200 h. **(G’,H’)** Magnified image of the circular and longitudinal smooth muscle. **(I-J)** Quantification of longitudinal **(I)** and circular **(J)** smooth muscle volume (n=3 mice/group). Superovulation results in increased longitudinal and circular smooth muscle volume compared to control; and imatinib treatment rescues longitudinal smooth muscle volume in superovulated uteri (*p* < 0.10). **(K-L)** Spatiotemporal plots (myographs) reflecting smooth muscle contractility at GD3 1200 h in superovulation+vehicle **(K)** and superovulation+imatinib **(L)** uteri. **(M-O)** Contraction metrics: **(M)** amplitude, **(N)** frequency, and **(O)** velocity in superovulation + vehicle/imatinib (vehicle n=2 mice; imatinib n=3 mice). White lines on the myographs indicate example waves. Amplitude is reduced in superovulation+imatinib treated mice, whereas frequency and velocity remain unchanged compared to superovulated mice. Statistical test: Mann–Whitney U-test. ns, non-significant. Red dashed lines indicate the median. M, mesometrial pole; AM, anti-mesometrial pole. CON, control; SO, superovulation; SO+VEH Ima, vehicle for Imatinib; SO+Ima, superovulated mice treated with Imatinib; SO+VEH Avap, vehicle for Avapritinib; SO+Avap, superovulated mice treated with Avapritinib; CM, circular smooth muscle; LM, longitudinal smooth muscle. <u>Scale bars:</u> **B-E:** 200 µm; inserts in **B, E:** 50 µm, **C,D:** 100 µm; **G,H** - IF: 200 µm; **G’,H’** – magnified image: 50 µm.

Imatinib has been implicated in relaxing intestinal smooth muscle^58^, thus, we evaluated the impact of imatinib treatment on uterine smooth muscle architecture and contractile function (**Fig. 7H,H’**). Quantitative analysis revealed that normalized longitudinal smooth muscle volume was rescued in superovulated mice treated with imatinib, reaching levels comparable to controls (Median: control=0.255 nu, Superovulation+Imatinib= 0.281 nu) (**Fig. 7I**). In contrast, imatinib treatment in superovulated mice did not rescue circular smooth muscle volume (n=4 mice per group; **Fig. 7J**) or excess collagen deposition (**Supplementary Fig. 12**). For contractile function, imatinib treatment of superovulated uteri reduced contraction amplitude (**Supplementary Video 3 and 4, Fig. 7M**, median: control = 0.1714 mm, Superovulation+vehicle = 0.2052 mm, Superovulation+Imatinib = 0.1308 mm). Contractile frequency and velocity were not rescued with imatinib treatment of superovulated uteri (**Fig. 7N, O**).

We next determined whether PDGFRA inhibition of superovulated uteri would also rescue post-implantation chamber development. At GD4 1800 h, since there were no longitudinal folds in superovulated uteri treated with imatinib, no embryo trapping was observed. Control uteri displayed the expected elongated V-shaped implantation chambers (n=4 mice; **Fig. 8A**), and imatinib-treated superovulated mice exhibited embryos at the AM pole of the luminal epithelium in significantly smaller implantation chambers resembling an earlier GD4 0000h time point (n=7 mice; **Fig. 8B, D**^41^). 18 hours later, at GD5 1200 h, implantation chambers in imatinib-treated superovulated mice had progressed to fully formed V-shaped chambers (n=3 mice; **Fig. 8C**), with embryo-uterine alignment restored (**Fig. 8D**; median angle: 88° at GD4 1800 h; 30.6° at GD5 1200 h). Again, no embryo trapping was observed. By mid-gestation, due to imatinib treatment in superovulated uteri, there was a moderate increase in embryo viability as determined by the presence of a strong heartbeat. At GD13 1200h, 69.76% of embryos in imatinib-treated superovulated mice were viable, whereas in superovulation, 59.45% of embryos were viable (**Supplementary Fig. 13**). Although this difference did not reach statistical significance, the trend suggests that PDGFRA inhibition may partially restore embryo survival under superovulatory conditions. Together, these findings demonstrate that PDGFRA inhibition after ovarian stimulation restored key superovulatory phenotypes, including longitudinal to transverse fold transition, longitudinal smooth muscle volume, contraction amplitude, and embryo implantation chamber formation, supporting a central role for PDGFRA signaling in superovulation-induced uterine aberrations during the pre-implantation period.

**Figure 8.**
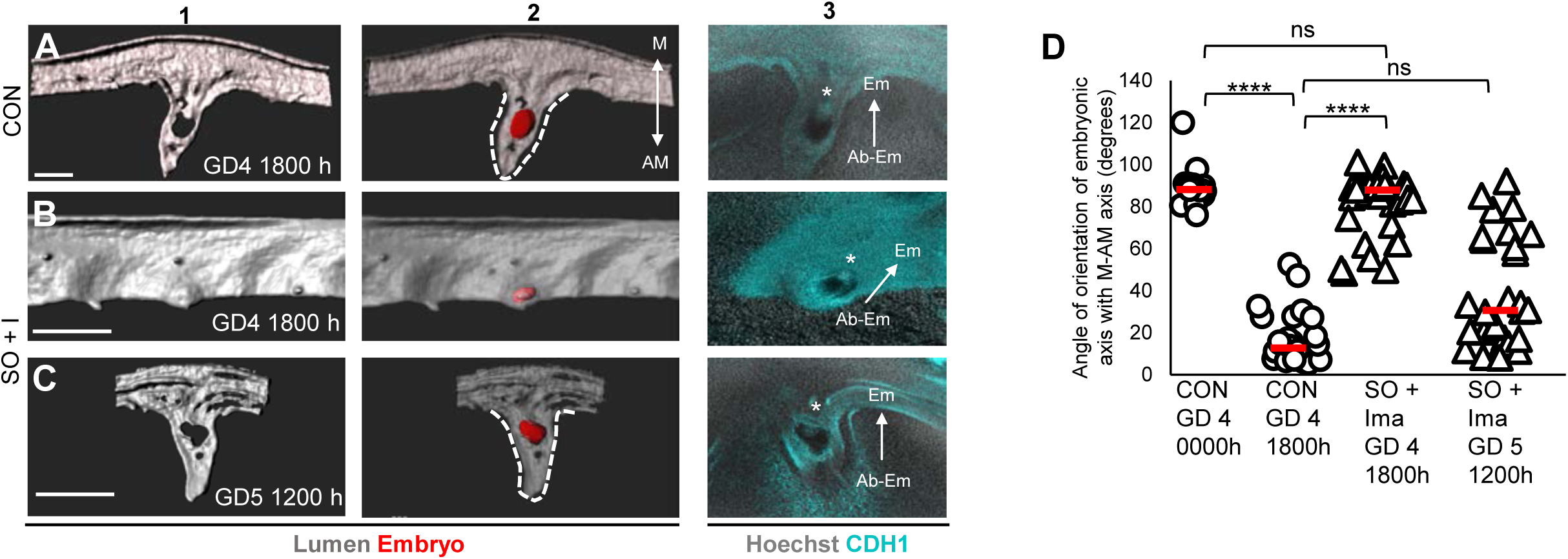
PDGFRA inhibition prevents embryo trapping in superovulated mice. **(A-C)** 3D surface and optical slice views of implantation sites at GD4 1800 h in control **(A)** and superovulation+imatinib uteri **(B)**, and at GD5 1200 h in superovulation+imatinib uteri **(C)**. Superovulation + imatinib uteri exhibit smaller implantation chambers at GD4 1800h **(B)** compared with control **(A)**. By GD5 1200 h, chamber length in superovulation+imatinib uteri **(C)** is comparable to control uteri at GD4 1800 h **(A)**, indicating that embryos are not trapped as observed in superovulated uteri but there is a delay in implantation chamber formation (Fig. 2F). Panel 1: 3D lumen surface (gray). Panel 2: transparent 3D lumen and embryo surface (red). Panel 3: frontal optical slice view. Asterisks: inner cell mass. **(D)** Quantification of embryo orientation relative to the M-AM axis in control GD4 0000 h (n = 3 mice, n_e_=15 embryos), control GD4 1800 h (n = 4 mice, n_e_=32 embryos), superovulation+imatinib GD4 1800 h (n = 3 mice, n_e_=35 embryos), and superovulation+imatinib GD5 1200 h (n = 3 mice, n_e_=28 embryos). Statistical test: *p* < 0.05, Mann–Whitney U-test; ns, non-significant. Red dashed lines indicate the median angle. M, mesometrial pole; AM, anti-mesometrial pole; Em, embryonic pole; Ab-Em, abembryonic pole; n_e_, number of embryos. CON, control; SO+I, superovulated mice treated with Imatinib. Scale bars: **A:** 300 μm; **B:** 400 μm; **C:** 500 μm.

## DISCUSSION

Ovarian stimulation is widely used in assisted reproductive technologies (ART), yet how it impacts the uterine milieu, and subsequent embryo implantation remains poorly understood. Using a mouse model, we demonstrate that ovarian stimulation disrupts remodeling of luminal epithelial folds, induces aberrant smooth muscle structure and contractility, results in increased collagen deposition, alters stroma-immune communication and elevates PDGFRA levels in the stroma and smooth muscle compartments. The heightened PDGFRA levels are also observed in endometrial biopsies from women in stimulated cycles. In the mouse, the structural and molecular alterations due to ovarian stimulation lead to impaired implantation chamber formation, embryo misalignment, and post-implantation embryo loss. Importantly, these defects can be reversed either by a rest period with normal estrus cycling or by pharmacological inhibition of PDGFRA signaling in the stimulation cycle. Using ovarian stimulation, our study identifies novel uterine structural and molecular targets that impact post-implantation embryo development. These targets can be modulated to improve pregnancy outcomes in clinical conditions associated with implantation failure.

In mouse models, it has been demonstrated that both embryo quality and the uterine environment are affected by ovarian stimulation^29,44,59^. Further, elevated levels of estrogen and progesterone during mouse ovarian stimulation have been shown to compromise both pre-and post-implantation embryo quality^14,29,44,59,60^ with dying embryos observed at cleavage stage, around the time of implantation, at mid-pregnancy, and near parturition^61–63^. Importantly, poor quality embryos observed post-fertilization typically undergo attrition during the pre-implantation period, such that morphologically healthy embryos surviving to blastocyst stage are generally considered capable of implantation^64^. In agreement with this, when healthy blastocysts from control and SO mice are transferred into control pseudopregnant uteri, embryo survival at mid-gestation or birth was not significantly different^30^. On the other hand, when healthy blastocysts are transferred into SO pseudopregnant uteri only 11% of the implanted embryos result in live pups compared to 66% in the control pseudopregnant uteri, suggesting a defect in the uterine milieu of the SO mouse^30^. Since we observe morphologically healthy blastocysts during pre-implantation stages after ovarian stimulation, we posit that embryo quality is not the primary driver of the post-implantation embryo loss in our studies. Our study suggests that these losses are primarily due to implantation chamber defects arising right after implantation, with embryo loss occurring consequently between GD5 and mid-gestation. The earliest phenotype we identify in the ovarian stimulated mice is failure of the uterine luminal epithelium to undergo transition from longitudinal to transverse folds, which results in embryo trapping and defective implantation chamber formation. These findings are consistent with other mouse models where loss of WNT5A or RBPJ resulted in aberrant folding, embryo trapping and consequent loss of embryos at mid-gestation^41,43^. Ovarian stimulated mice also display disorganization of uterine smooth muscle fibers, enlarged smooth muscle cells in both the longitudinal and circular muscle layers and aberrant uterine contractility during the pre-implantation stages, suggesting structural and functional disruptions in the smooth muscle compartment.

Ovarian stimulation induced abnormal peri-implantation uterine environment in fresh embryo transfer cycles is linked to increased adverse perinatal outcomes when compared to frozen embryo transfer cycles^22^. Our study shows that a rest and recovery period fully restored epithelial fold morphology, implantation chamber formation, and mid-gestation pregnancy outcomes. These findings provide a mechanistic basis for the improved implantation outcomes observed with frozen embryo transfer cycles. Collectively, our data indicate that the disruption in uterine architecture caused by a single cycle of ovarian stimulation is transient and reversible and explains why deferring transfer may enhance uterine receptivity and embryo implantation rates. Ovarian stimulation in women leads to elevated numbers of B cells, effector CD4+ T-cells, and macrophages and a reduction in NK cell numbers and alteration of immune cell migration pathways^17,65^. Using transcriptomics studies in mice have shown pre-implantation enrichment of immune pathway related genes and a reduction in B cells and NK cells in ovarian stimulated uteri^33,66^. In agreement, our study shows immune cell perturbations in stimulated uteri with an increase in T-cells and dendritic cells and perturbed NK cell gene signatures. Further, both ovarian stimulated deep and superficial stromal compartments displayed increased outgoing collagen based signaling with the T cells and dendritic cells suggesting a disruption of the ECM under ovarian stimulation conditions. Our study and previous studies suggest that genes involved in endometrial remodeling, including extracellular matrix genes, gap junction genes, tight junction genes and adherens junction genes are dysregulated during ovarian stimulation^33^. In humans, transcriptomic profiling of the endometrium following ovarian stimulation also shows differential expression of extracellular matrix remodeling protease coding genes, including *MMP10*, *HPSE*, *MMP2* and *TIMP1*, suggesting that ovarian stimulation disrupts the proteolytic balance required for normal ECM remodeling^19,23^. We also observed an enrichment of ECM-related GO terms in the ovarian stimulated stromal compartment alongside transcriptional upregulation of pathways associated with ECM organization and excess fibrillar collagen deposition in both the stroma and smooth muscle compartments. Stromal fibroblast-derived ECM remodeling is a well-established regulator of epithelial architecture and function across multiple organs, including the mammary gland^67,68^ and the embryonic gut^69^ and may contribute to aberrations in epithelial folding observed in our stimulated uteri. Further, dynamic regulation of fibrillar collagens during peri-implantation uterine remodeling is well established in both humans and mice^70,71^. Importantly, fibrillar collagen tension in the endometrium has been directly linked to embryo implantation rates, and enzymatic reduction of collagen improves implantation in mouse models^72^. Similar to ovarian stimulated mouse uteri in our study, aberrant collagen deposition and fibrosis have been observed in Asherman’s syndrome^73^ and continuously cycling aging mice^74^, both conditions linked to a decline in fertility. Thus, the heightened immune engagement and a pro-inflammatory and high collagen pro-fibrotic state induced by ovarian stimulation may be important contributors to the implantation defects observed under conditions of ovarian stimulation.

PDGFRA signaling promotes collagen deposition and ECM remodeling in connective tissues^75^ and we found elevated pre-implantation PDGFRA expression in stromal and smooth muscle compartments of ovarian stimulated uteri. Additionally, PDGFRA-mediated epithelial-mesenchymal interactions are known to guide epithelial folding and alveolar septation in the lung^76^ and aggregation of PDGFRA-high sub-epithelial stromal cells in the intestine drives villus (epithelial fold) formation^55^. These data are consistent with a role for elevated PDGFRA signaling in dysregulated epithelial folding in ovarian stimulated uteri. Further, PDGFRA signaling is essential for proper smooth muscle layer differentiation^77,78^. The enhanced smooth muscle contraction amplitude observed with elevated PDGFRA expression in the ovarian stimulated mouse uteri likely involve multiple downstream mechanisms. PDGFRA signaling controls SRF nuclear localization through activation of RAC1 and regulation of actin dynamics, both of which are central to contractile gene expression^79^. Additionally, PDGFRA activation triggers downstream PI3K/AKT, MAPK/ERK, and PLCγ/PKC pathways that collectively contribute to enhanced contractility^79,80^. The PLCγ pathway is particularly relevant in this context, as it initiates calcium mobilization directly coupled to smooth muscle contraction^80^. Whether activation of these PDGFRA-mediated downstream signaling cascades contributes to the aberrant uterine contractility in ovarian stimulated conditions remains an important question for future investigation.

Pharmacological inhibition of excess PDGFRA signaling in ovarian stimulated mice rescued the epithelial fold transition, uterine contractility and implantation chamber formation in our study. Imatinib is a multi-tyrosine kinase inhibitor that inhibits PDGFRA kinase activity^81^ and has previously been shown to relax longitudinal smooth muscle by blocking signals upstream of excitation-contraction coupling^58^. Avapritinib is a selective PDGFRA inhibitor^82,83^. Although avapritinib adversely affected embryo morphology, both compounds rescued epithelial folding and imatinib also rescued uterine contractility defects in the ovarian stimulated mouse uteri. These findings establish PDGFRA as a critical regulator of uterine tissue folding and smooth muscle contractility. PDGFRA inhibition rescued structural defects and implantation chamber formation and mildly improved mid-gestation pregnancy outcomes in the ovarian stimulated mice. Importantly, we found PDGFRA expression to be elevated in uterine biopsies from women undergoing ovarian stimulation, suggesting that the mechanisms identified in our mouse model may be relevant to human infertility. Elevated PDGFRA expression coincided with rising progesterone levels; however PGR binding sites were not observed in the *Pdgfra* promoter region^84^. Whether PDGFRA expression in uterine stromal cells is indirectly regulated by progesterone signaling, remains an important question for future investigation. Both imatinib and avapritinib are FDA-approved inhibitors currently used in oncological settings, and our evidence suggests that a short course of low-dose treatment may improve uterine epithelial architecture for conditions with poor implantation outcomes.

In summary, ovarian stimulation disrupts uterine epithelial folding, smooth muscle contractility, extracellular matrix composition including collagen deposition, and gene expression including stromal and smooth muscle PDGFRA signaling. These structural and molecular changes converge to impair implantation chamber formation, resulting in post-implantation pregnancy loss. The structural defects induced by ovarian stimulation were reversible by normal cycling or controlled short-term PDGFRA inhibition. Our study identifies PDGFRA as a key modulator of uterine contractility and non-cell autonomous regulation of epithelial folding, both of which are critical for implantation. Our research highlights novel structural and molecular pathways that are critical for implantation and can be leveraged to improve pregnancy rates in the case of assisted reproduction and recurrent implantation failure.

## MATERIALS AND METHODS

### Animals and treatments

All mouse studies were approved by the Michigan State University Institutional Animal Care and Use Committee. C57BL/6 mice of ages 6-10 weeks were used for the study. Superovulation (SO) was induced by injecting C57BL/6 female mice with 5 IU Pregnant Mare Serum Gonadotropin (PMSG) and 48 hours later with 5 IU Human Chorionic Gonadotropin (hCG). For SO followed by 4 weeks of rest (SO+4w rest), the female mice were first superovulated and mated with a vasectomized male to complete the hormonal cascade of SO. Mice were allowed a 4 week recovery period (∼5-6 estrus cycles) before mating. The female mice were mated with fertile males (for pregnancy time points) or vasectomized males (to induce pseudopregnancy). The appearance of a vaginal plug confirmed mating and was identified as gestational day (GD) 0.5. Mice were humanely euthanized, and uteri were dissected at GD1 1200 h, GD3 1200 h, GD4 1800 h, GD5 1200h, and GD13 1200 h.

For PDGFRA inhibition, C57BL/6 female mice were subjected to SO and mated as described earlier and injected intraperitoneally with 2 mg/mouse imatinib mesylate (SML1027, Sigma) or DI water (vehicle for Imatinib) or 0.187 mg/mouse avapritinib (#50-193-2168, Medchemexpress) or 0.94% DMSO+saline (vehicle for avapritinib) at GD2 2200h (10pm, before the transition of folding started). The mice were humanely euthanized, and uteri were dissected at GD3 1200 h, GD4 1800 h, GD5 1200h, and GD13 1200 h.

### Human endometrial biopsy samples

Using an Institutional Review Board (IRB)-approved protocol at Rutgers New Jersey Medical School and a university-affiliated fertility clinic, human endometrial biopsy samples were obtained from a cohort study of women in natural menstrual cycles (LH-triggered) and ovarian stimulation cycles (hCG-triggered), as described in Chemerinski et al^17^. Briefly, biopsies were collected at the periovulatory and mid-secretory phases of the menstrual cycle from women with regular ovulatory cycles and from women undergoing ovarian stimulation as part of artificial reproductive technology treatment. Tissue sections were fixed in 4% PFA overnight and processed for cryoembedding.

### Whole-mount immunofluorescence staining

Whole-mount staining was performed as previously described (Arora et al., 2016). Dissected uteri were fixed in DMSO:methanol (1:4) and stored at −20°C. The uteri were rehydrated in methanol:PBT (1% Triton X-100 in PBS) (1:1) for 15 min, followed by a PBT wash for 15 min. Samples were incubated in blocking solution (2% powdered milk in PBT) for 1 hour at room temperature. Uteri were incubated with primary antibodies at a 1:500 dilution in blocking solution for 5 nights at 4°C. Following incubation, uteri were washed with PBT 2 x 15 min and 4 x 45 min at room temperature, then incubated with secondary antibodies at 4°C for 3 nights. Uteri were washed with PBT 1 x 15 min and 3 x 45 min, dehydrated in methanol for 15 min, and bleached in 3% H₂O₂ in methanol overnight at 4°C. Samples were then washed in 100% methanol 2 x 15 min and 1 x 60 min and cleared in BABB (benzyl alcohol:benzyl benzoate, 1:2; Sigma-Aldrich, 108006, B6630). Primary antibodies used include rat anti-CDH1 (M108, Takara Biosciences) and rabbit anti-FOXA2 (ab108422, Abcam). Secondary antibodies used include Alexa Fluor-conjugated donkey anti-rabbit 555 (A31572, Invitrogen) and goat anti-rat 633 (A21094, Invitrogen), with Hoechst (Sigma-Aldrich, B2261) as a nuclear marker.

### Cryoembedding and sectioning

Uterine tissue samples or endometrial biopsies were fixed in 4% PFA overnight at 4°C. Samples were washed in PBS for 3 x 5 min at room temperature, then serially transferred through 10%, 20%, and 30% sucrose solutions at 4°C. Samples were placed in cryomolds, embedded in OCT, and frozen at −80°C. Frozen blocks were sectioned at 7 μm thickness using a cryostat and mounted onto glass slides, which were stored at −80°C until use.

### Cryosection immunofluorescence staining

Cryosection slides were thawed at room temperature and outlined with a hydrophobic barrier. Sections were washed in PBS for 5 min and incubated in blocking solution (0.1% powdered milk in PBT) for 20 min at room temperature.

For sections processed for confocal imaging, a two-day staining protocol was used. Sections were incubated with primary antibodies diluted in blocking solution overnight at 4°C, followed by PBS washes for 3 x 5 min. Sections were then incubated with secondary antibodies diluted in 0.1% PBT for 1 hour at room temperature, washed again with PBS for 3 x 5 min, and mounted for imaging. Primary antibodies used for mouse uterus sections were rat anti-CDH1 (M108, Takara Biosciences; 1:500) and goat anti-PDGFRA (AF1062, R&D Systems; 1:200). Alexa Fluor-conjugated secondary antibodies used for mouse sections were donkey anti-goat 555 (A21432, Invitrogen; 1:500) and goat anti-rat 488 (A21208, Invitrogen; 1:500). Primary antibodies used for human uterine biopsy sections were mouse anti-CDH1 (ab1416, Abcam; 1:200) and rabbit anti-PDGFRA (ab203491, Abcam; 1:200). Alexa Fluor-conjugated secondary antibodies used for human sections were donkey anti-mouse 647 (A31571, Invitrogen; 1:500) and donkey anti-rabbit 555 (A31572, Invitrogen; 1:500). Nuclei were counterstained with Hoechst (B2261, Sigma-Aldrich; 1:500) for all sections.

For sections processed for collagen detection using second harmonic generation (SHG) imaging, a one-day staining protocol was used. Following blocking, sections were incubated for 2 hours with mouse anti-ACTA2-FITC (F3777, Sigma-Aldrich; 1:250), washed with PBS for 3 x 5 min, and mounted for imaging.

### Confocal microscopy

Whole tissue samples were imaged using Leica TCS SP8 X Confocal Laser Scanning Microscope System with white-light laser and 10X air objective. The entire length and thickness of the uterine horn was imaged using the tile scan function with z-stacks of 7μm. Images were merged using Leica software LASX version 3.5.7. Tissue sections were imaged using Leica TCS SP8 X Confocal Laser Scanning Microscope System with white-light laser and 20X water immersion objective.

### Second Harmonic Generation (SHG) imaging

Leica SP8 DIVE laser multiphoton microscope equipped with Spectra-Physics Insight X3 dual beam (680 to 1,300 nm tunable and 1,040 nm fixed) and 4Tune, tunable, super sensitive hybrid detectors (HyDs) with a 25x objective lens was used to acquire images of the stroma and smooth muscle in a transverse section (LAS X software version 3.5.7.23225). A wavelength of 950 nm was used to detect a collagen signal at 475 nm. The acquired images were subsequently analyzed using Imaris v9.2.1. (Bitplane; Oxford Instruments, Abingdon, UK)

### 3D reconstruction and image analysis

Image analysis was performed using Imaris v9.2.1 (Bitplane). The confocal image.LIF files were imported into the Surpass mode of Imaris. Using the channel arithmetic function, the FOXA2 signal of glands was subtracted from the epithelial CDH1 signal to isolate the lumen-only signal. The Surface module was then used to reconstruct the 3D surface of the lumen from the lumen-only channel. Embryo surfaces were reconstructed using the Manual mode of the Surface module from the Hoechst signal. Quantification of luminal folding angle (at GD3 1200h), embryo-uterine orientation (GD4 1800h and GD5 1200h), and the space between embryo and the decidua (GD5 1200h) was performed using the Measurement Points module in Imaris^41^.

### PDGFRA signal quantification

To quantify PDGFRA signal intensity from immunofluorescence-stained cryosections, images were analyzed using the Surface module in Imaris. Surfaces were created for nuclei (using Hoechst signal) and for PDGFRA signal within the stromal and smooth muscle compartments separately. Nuclear volume was used for normalization. Two metrics were determined: the total volume of stromal and smooth muscle compartments expressing PDGFRA in mouse tissue, and the volume of these compartments expressing low and high levels of PDGFRA based on fluorescence intensity. PDGFRA signal was classified into two intensity categories using pixel intensity thresholds: in mouse tissue, low intensity was defined as < 200 IU and high intensity as > 200 IU, while in human tissue, low intensity was defined as < 128 IU and high intensity as > 128 IU. The volume of voxels within each compartment and intensity category was extracted using the volumetric function in Imaris and exported to Microsoft Excel, where the volume of low- and high-intensity PDGFRA signal was normalized to the nuclear volume of the respective compartments.

### Spatiotemporal mapping protocol

A spatiotemporal mapping protocol to quantify uterine contractions waveform metrics was used^46^. C57Bl6 mice carrying the *Gt(ROSA)26Sor^tm4(ACTB-tdTomato,-EGFP)Luo^*/J allele (Jackson labs #007676) and expressing membrane Tomato transgene in all cells was used^85^. Briefly, following humane euthanasia and within 5 min of dissection, the uterus was placed in Krebs buffer (Zen bio, cat# KRB-1000/ Z990181) under a Leica MZ10F fluorescence stereo microscope with a fluorescence filter for Texas Red. Using LAS X software, time-lapse uterine contractions were recorded with images captured every 200 ms for 5 min (total 1500 frames). Contraction videos were post-processed and subjected to image analysis using a python code^46^. Spatiotemporal myographs were extracted based on fluorescence intensity and waves were manually traced on these myographs to calculate contraction wave metrics. Amplitude was used to derive contraction strength, velocity quantified contraction speed, and frequency measured the number of contractions over a set time period.

### Single cell isolation

Uterine horns from SO and control mice (n = 2 mice per group) were dissected at GD3 0900 h in sterile phenol red-free 1× HBSS with Ca and Mg (Gibco, cat# 14025092) and placed on ice. Tissues were minced with scissors and collected in gentleMACS C-tubes (Miltenyi Biotec, cat# 130-093-237) containing the Multi Tissue Dissociation Kit 2 enzyme cocktail (Miltenyi Biotec, cat# 130-110-203). The C-tubes containing the minced tissue and enzyme cocktail were placed in the Gentle MACS nutator inside an incubator at 37°C for enzymatic dissociation for 30 minutes. After this incubation period, samples were transferred to the Gentle MACS Dissociator for mechanical dissociation, where the m_intestine_01 program was run. The sample was again incubated at 37°C for another 30 minutes followed by another mechanical dissociation step with the m_intestine_01 program. The sample was then incubated at 37°C for another 15 minutes followed by the m_heart_01 program. Digested tissues were strained through a 40 µm nylon cell strainer (Fisher Scientific, cat# 22-363-547) into a 50 mL Falcon tube and washed with 10 mL of MACS buffer. The flow-through and filtrate were centrifuged at 2500 rpm for 5 min at room temperature and the supernatant was removed. To remove red blood cells, 10 mL of 1× Red Cell Lysis Buffer (Miltenyi Biotec, cat# 130-094-183) was added to the cell pellets, which were gently dissociated and incubated for 3 min at room temperature, followed by centrifugation at 2500 rpm for 5 min. Dead cell removal was performed using the MACS Dead Cell Removal Kit (Miltenyi Biotec, cat# 130-090-101). Cells were resuspended in 100 µL of Dead Cell Removal Microbeads per 10⁷ cells and incubated at room temperature for 15 min. MS columns were placed in the magnetic field of a MACS separator and rinsed with 500 µL of 1× binding buffer. One milliliter of binding buffer was added to the cell-bead suspension and the mixture was loaded onto the column. Columns were rinsed four times with 500 µL of 1× binding buffer, and the effluent containing the live cells was collected in 5 mL round-bottom Falcon tubes. The live cell fraction was centrifuged at 300 × g for 5 min and resuspended in 1ml of cold MACS buffer.

### RNA isolation, library preparation and sequencing

Libraries were generated and sequenced by the Van Andel Institute Genomics Core (Grand Rapids, MI). Cells were processed using the 10× Chromium Next GEM Single Cell 5′ GEM kit v2 (10× Genomics, Pleasanton, CA) according to the manufacturer’s instructions, targeting an output of 10,000 cells per sample using a 10× Genomics Chromium Controller. Single-cell suspensions in PBS with 0.04% BSA were assessed for quantity and viability using the CytoFLEX S (Beckman Coulter, Indianapolis, IN), and 16,000 cells per sample were loaded onto the Chromium Controller. Single cells were captured in gel beads in emulsion (GEMs), where they were lysed and the released RNA was barcoded and converted to cDNA. Quality and quantity of the finished gene expression libraries were assessed using an Agilent DNA High Sensitivity chip (Agilent Technologies) and the QuantiFluor dsDNA System (Promega, Madison, WI). 2 × 100 bp paired-end sequencing was performed on an Illumina NovaSeq 6000 sequencer (Illumina, San Diego, CA) using an S4 300-cycle sequencing kit (v1.5) to a minimum depth of 20,000 reads per cell. Base calling was performed using Illumina RTA3, and sequencing output was demultiplexed and converted to FastQ format using Cell Ranger (10× Genomics, v3.1.0, RRID:SCR_017344). Raw data has been deposited at the NCBI Gene Expression Omnibus.

### Single-cell RNA-sequencing analysis

For scRNA-seq, demultiplexed sequencing reads were processed and aligned to the Mus musculus genome assembly GRCm39 (mm39) using STAR (v2.7.9a, RRID:SCR_004463) with 10× Genomics Cell Ranger. Samples were merged using the integration anchors function of the Seurat package (v5.1.0, RRID:SCR_016341) in R. Genes expressed in fewer than three cells were excluded, as were cells expressing fewer than 200 genes or mitochondrial gene content > 5% of total unique molecular identifier counts. Data were normalized using a global-scaling normalization method, whereby feature expression measurements for each cell were divided by total expression, multiplied by a scale factor of 10,000, and log-transformed. The top 2,000 most variable genes were identified using the FindVariableFeatures function and normalized using the ScaleData function. Based on an elbow plot generated using the ElbowPlot function, 25 principal components were selected for downstream analyses. Cell clusters were identified using the FindNeighbors and FindClusters functions. For visualization, UMAPs were generated using the RunUMAP, FeaturePlot, and DimPlot functions. Dot plots illustrating gene expression across clusters and violin plots showing expression distributions of genes of interest were generated using the DotPlot and VlnPlot functions, respectively. Bar plots were generated using the dittoSeq R package (v1.16).

### Differential gene expression visualization, gene ontology analysis, and CellChat

Differentially expressed genes (DEGs) identified from each cell cluster (luminal epithelial, stromal, smooth muscle, and immune cells) were visualized using volcano plots generated with the EnhancedVolcano package (v1.22.0) in R. Genes with an adjusted p-value < 0.05 and an absolute log2 fold change > 0.2 were considered statistically significant. Volcano plots were constructed using the average log2 fold change (avg_log2FC) on the x-axis and the negative log10 of the adjusted p-value on the y-axis, with x-axis limits adjusted per cluster. Genes that were significantly different are highlighted in red and those that are not different are in blue text. Select genes of biological interest were highlighted with labels and connector lines. Gene ontology analysis was determined with clusterProfiler package (version 4.20.0) using MSigDB (PMID: 16199517) reference databases. Ligand-receptor cellular communication analysis was determined with the CellChat (v2.1.2, RRID: SCR_021946) R package (PMID: 33597522, 39289562). Briefly, communication probabilities were calculated between cell types using the functions identifyOverExpressedGenes and identifyOverExpressedInteractions. For visualizing pathways overrepresented in cell-cell communication the CellChat function *netVisual_chord_gene* and *netVisual_bubble* was used.

### Human endometrial biopsy bulk RNA Sequencing Analysis

GEO Dataset GSE220044 that has bulk RNA-sequencing data for human endometrial biopsies was used^17^. Trimmed reads were mapped to Homo sapiens genome assembly GRCh38 (hg38) using STAR (v2.7.9a, RRID:SCR_004463). Reads overlapping Ensembl annotations (v110) were quantified with STAR prior to model-based differential expression analysis using the edgeR-robust method. For differential expression analysis, genes with low counts per million (CPM) were removed using the filterByExpr function from edgeR (RRID:SCR_012802). Genes were considered differentially expressed if the FDR corrected p-values were less than 0.05.

### Statistics

Statistical analysis was performed using GraphPad Prism (version 10.2.3). Data were assessed for normality and variance before selecting statistical tests. Unpaired Student’s t-test, Mann–Whitney U test, or Wilcoxon rank-sum test were used when comparing two groups. Kruskal–Wallis test or Welch’s t-test were used when comparing three or more groups. Dunn’s multiple comparison test was applied in conjunction with the Kruskal–Wallis test where appropriate. A *p*-value < 0.05 was considered statistically significant.

## ACKNOWLEDGEMENTS

We thank Sara Makaremi for help with SHG microscopy, Marie Adams for help with designing the single cell RNA sequencing experiment, Hannah Lufkin and Prof. Asgi Fazleabas for conceptual discussions and Kaylie Chiles for critical review of the manuscript.

## AUTHOR CONTRIBUTIONS

HRK, MM and RA conceptualized the study and designed the experiments. HRK, MM, LZ, CC, RY performed the experiments. HRK, MM, ENP, GWB, NCD and RA validated the data and performed the analyses. HRK, MM, LZ and RA prepared the figures and wrote and edited the manuscript. All authors reviewed and accepted the final version of the manuscript.

## DATA AVAILABILITY

All data supporting the findings of this study are available within the article or its supplementary data.

---

**Supplementary Figure 1.**
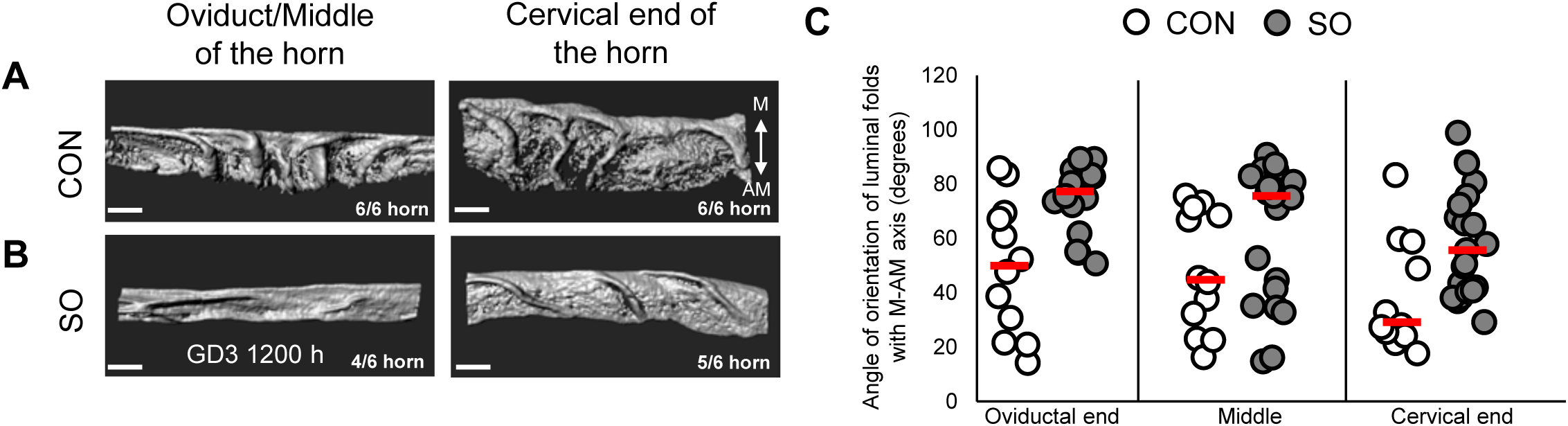
Longitudinal folds are observed in the oviductal and middle regions in superovulated uteri at GD3 1200h. **(A-B)** 3D reconstructions of control **(A)** and superovulated **(B)** uterine lumen demonstrate transverse folding throughout the uterine horn in control, whereas superovulated uteri display longitudinal folds in the oviductal and middle regions and transverse folding at the cervical end. **(C)** Quantification of fold angle relative to the M–AM axis, comparing the oviductal, middle, and cervical regions of control and superovulated uterus (n = 3 mice/group). CON, control; SO, superovulation. Scale bars: **A-B**: 300 µm.

**Supplementary Figure 2.**
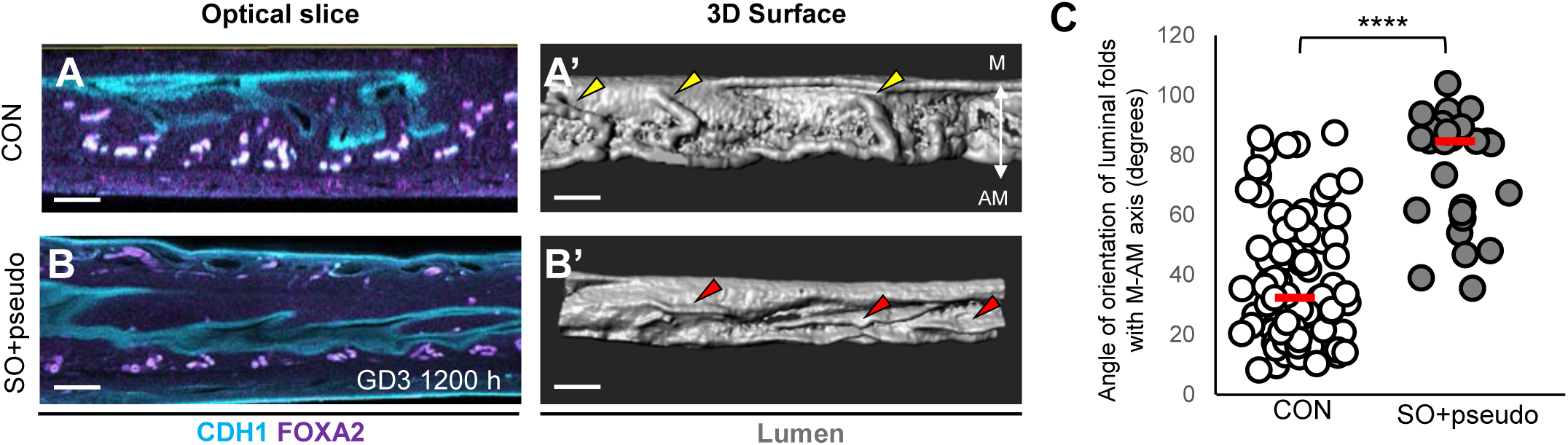
Failure of fold transition in superovulated mice occurs independently of embryos at GD3 1200h. **(A-B)** Optical confocal slices of control **(A)** and superovulation+pseudopregnant **(B)** uterus. **(A’-B’)** 3D reconstruction of the lumen in **A** and **B** demonstrates transverse folds (yellow arrowheads) in control and longitudinal folds (red arrowheads) in superovulation+pseudopregnant uterus. **(C)** Quantification of fold angle relative to the M–AM axis in control and superovulation+pseudopregnant uteri (n = 4 mice/group). Statistical test: *p* < 0.05, Mann–Whitney U-test. Red dashed lines indicate the median angle. M, mesometrial pole; AM, anti-mesometrial pole; CON, control; SO+pseudo, superovulated mice mated with a vasectomized male (pseudopregnant). <u>Scale bars:</u> **A,A’,B,B**’: 200 µm.

**Supplementary Figure 3.**
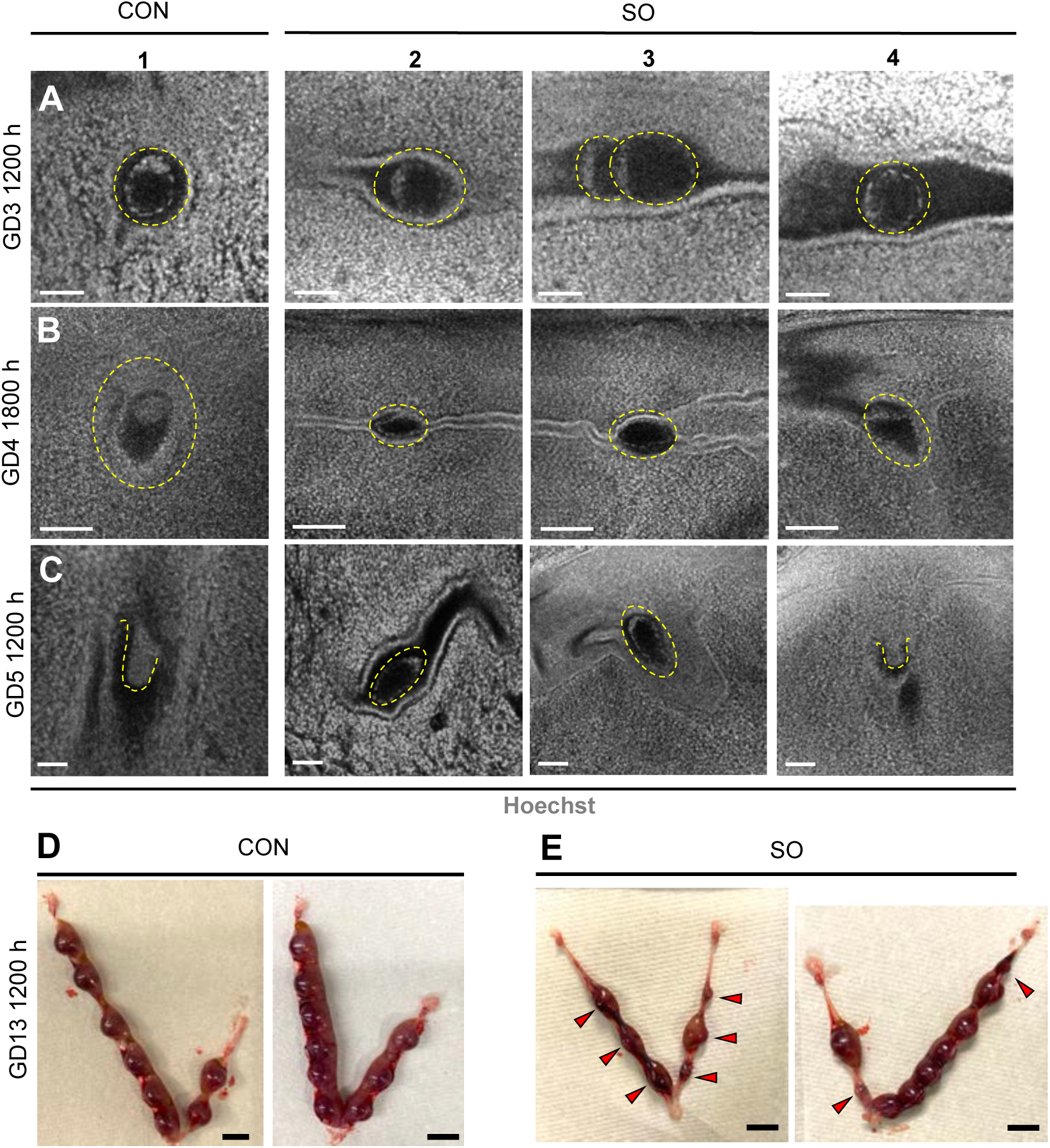
Superovulation compromises embryo morphology and post-implantation survival. **(A-C)** Optical slice images of embryos obtained by confocal imaging at GD3 1200h **(A)**, GD4 1800h **(B)**, and GD5 1200h **(C)**. Panel 1: control embryos, and Panels 2–4: superovulated embryos. Yellow dotted circles: embryo locations. At GD3 1200 h, all embryos in superovulated mice are at the blastocyst stage. At GD4 1800 h, superovulated embryos in panels 2 and 3 are trapped in a longitudinal fold and appear smaller compared to controls and superovulated embryos in panel 4 are from a normal V-shaped chamber. At GD5 1200 h, some superovulated embryos are morphologically deformed (panels 2 and 3) compared to controls, whereas superovulated embryos in Ch_norm_ appear similar to controls. **(D-E)** Uterine horns with embryos at GD13 1200 h in control **(D)** and superovulated **(E)** uteri (n=5 mice/group). Red arrowheads indicate resorption sites in SO. Ch_norm_, long chamber; CON, control; SO, superovulation. Scale bars: **A,C:** 50 µm; **B:** 100 µm; **D,E:** 5mm.

**Supplementary Figure 4.**
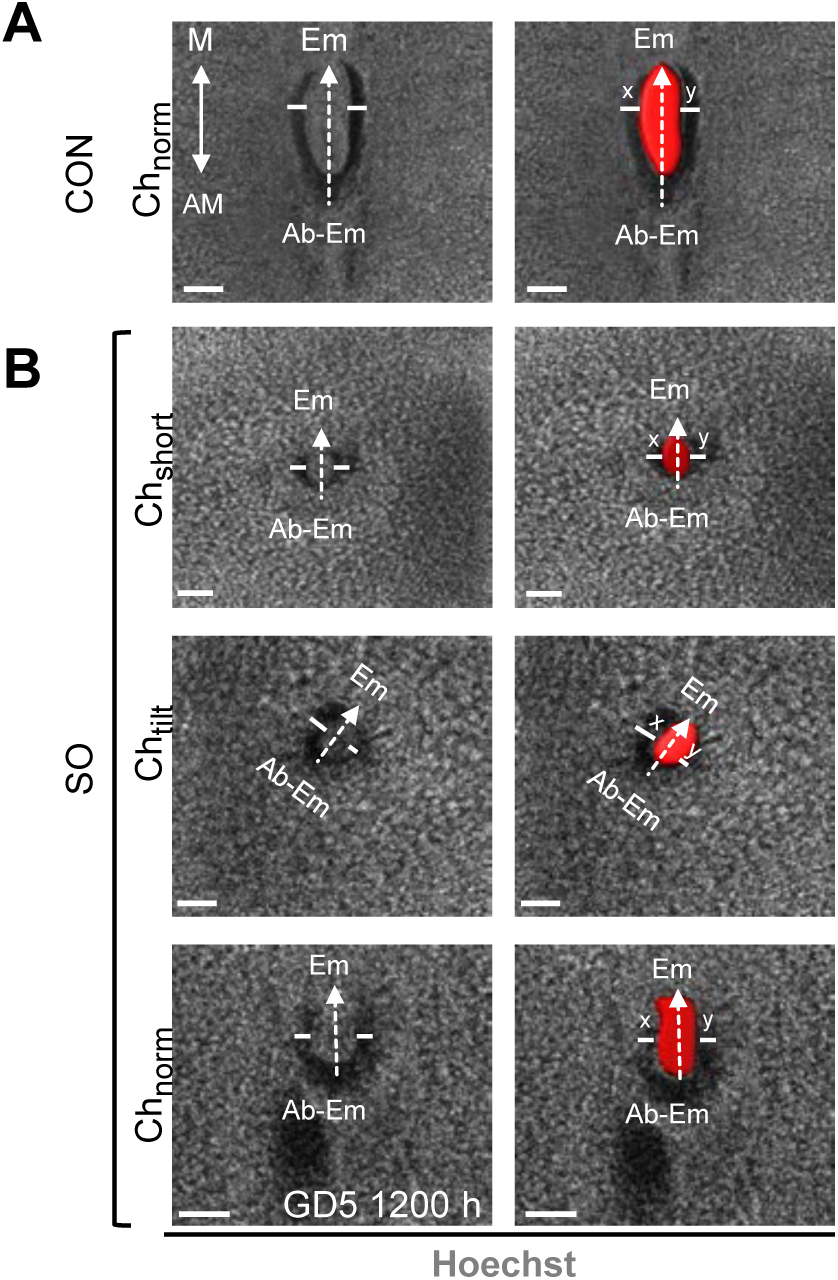
Embryo-uterine axis misalignment reduces epiblast-decidua buffer space in superovulated mice with tilted chambers at GD5 1200 h. **(A-B)** Optical slice images of implantation sites in control **(A)** and superovulated **(B)** uteri. Superovulated embryos with a short chamber (Ch_short_) or normal chamber (Ch_norm_) display equal buffer space between the epiblast and maternal decidua, whereas superovulated embryos with a tilted chamber (Ch_tilt_) display uneven buffer space between the epiblast and maternal decidua. CON, control; SO, superovulation. Scale bars: **A Ch_norm_, B Ch_tilt_:** 100 µm, **B Ch_short_:** 50 µm, **B Ch_norm_:** 70 µm.

**Supplementary Figure 5.**
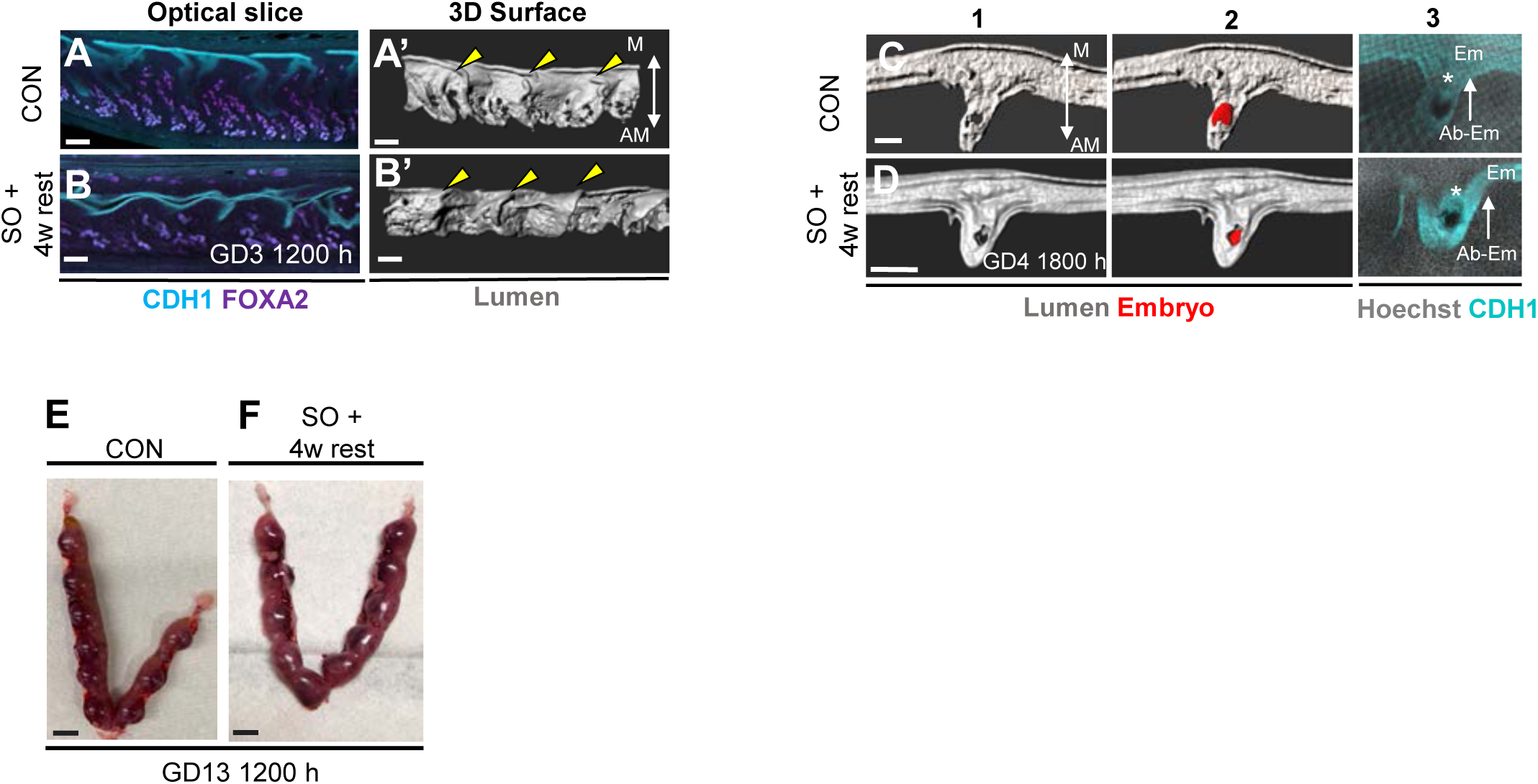
Resumption of normal estrous cycling in previously superovulated mice restores uterine architecture and implantation success. **(A-B)** Optical confocal slices of control **(A)** and superovulation+ 4-weeks rest **(B)** uteri at GD3 1200 h. **(A’-B’)** 3D reconstruction of the lumen in **A** and **B** (n=4 mice/group) demonstrates transverse folding (yellow arrowheads). **(C-D)** 3D surface and optical slice views of implantation sites at GD4 1800 h in control **(C)** and superovulation + 4-weeks rest **(D)** uteri (n = 4 mice/group). Asterisks: inner cell mass. **(E-F)** Uterine horns with embryos at GD13 1200 h in control (n=7 mice) **(E)** and superovulation + 4-weeks rest (n=6 mice) **(F)**. M, mesometrial pole; AM, anti-mesometrial pole. CON, control; SO + 4w rest, superovulation + 4-week rest. Scale bars: **A,B,C:** 200µm; **D:** 500 µm; **E,E’:** 5mm.

**Supplementary Figure 6.**
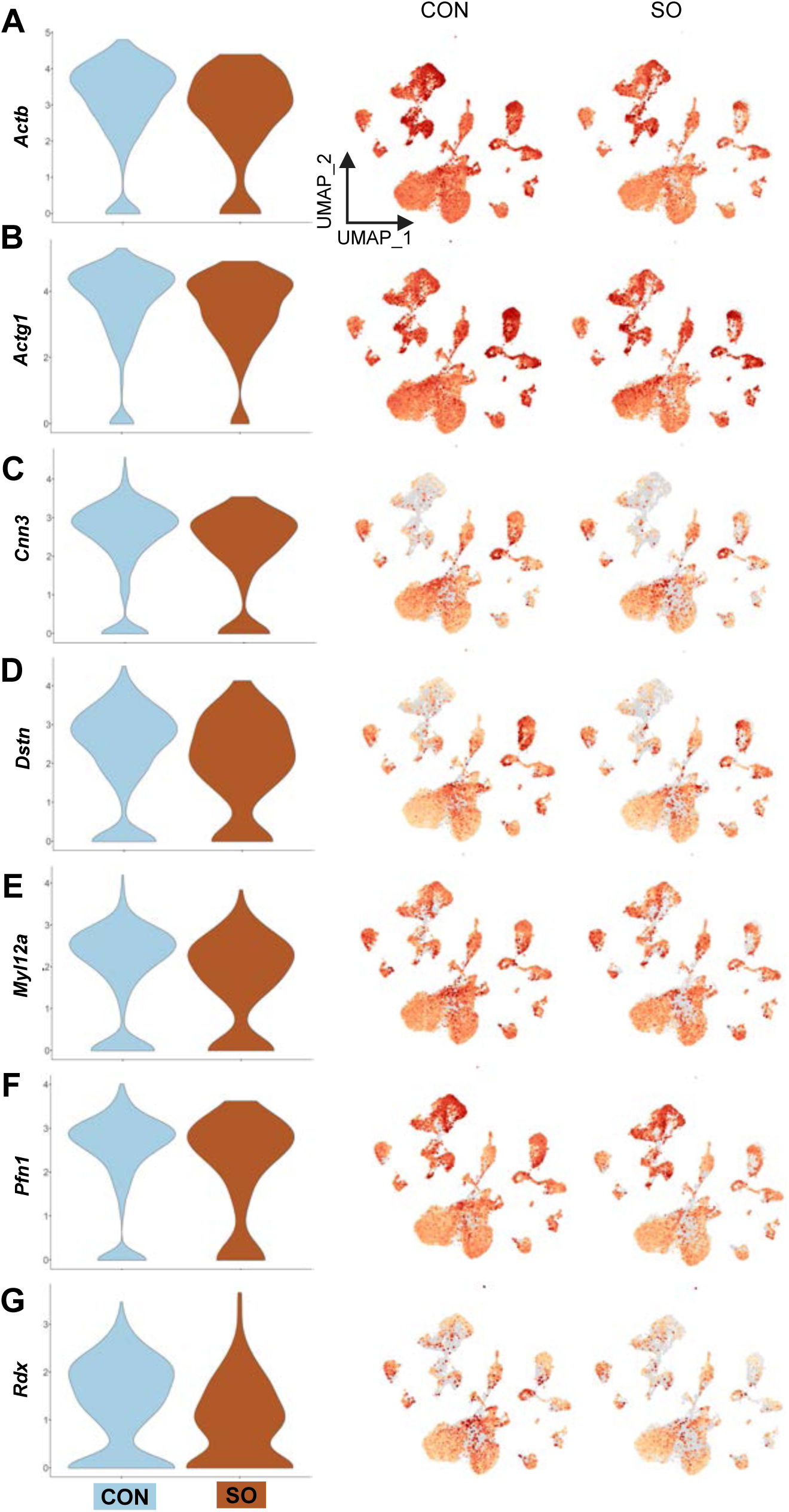
Downregulated luminal epithelial DEGs show broad expression across uterine cell types in superovulated mice. **(A-G)** Violin plots and UMAP depicting reduced expression of *Actb* **(A)**, *Actg1* **(B)**, *Cnn3* **(C)**, *Dstn* **(D)**, *Myl12a* **(E)**, *Pfn1* **(F)**, and *Rdx* **(G)** during superovulation compared to control. UMAP confirms that expression is broadly distributed across multiple uterine compartments and not specific to luminal epithelium. CON, control; SO, superovulation.

**Supplementary Figure 7.**
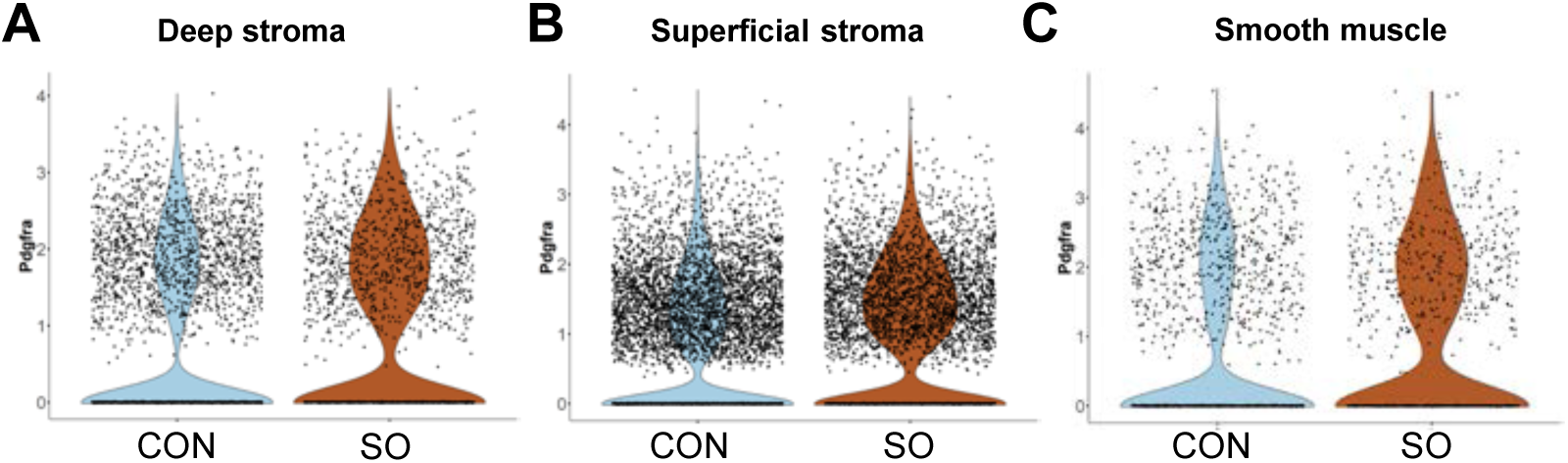
Elevated *Pdgfra* expression is observed in stroma and smooth muscle compartments of superovulated mice during pre-implantation. **(A-C)** Violin plots depicting elevated *Pdgfra* expression in deep **(A)**, superficial **(B)** stroma and smooth muscle **(C)** compartments during superovulation compared to control. Deep stroma (fold change = 1.31, FDR *p* = 3.45 × 10⁻¹²); superficial stroma (fold change = 1.30, FDR *p* = 2.26 × 10⁻³²); smooth muscle compartment (fold change = 1.35, FDR *p* = 0.001). CON, control; SO, superovulation.

**Supplementary Figure 8.**
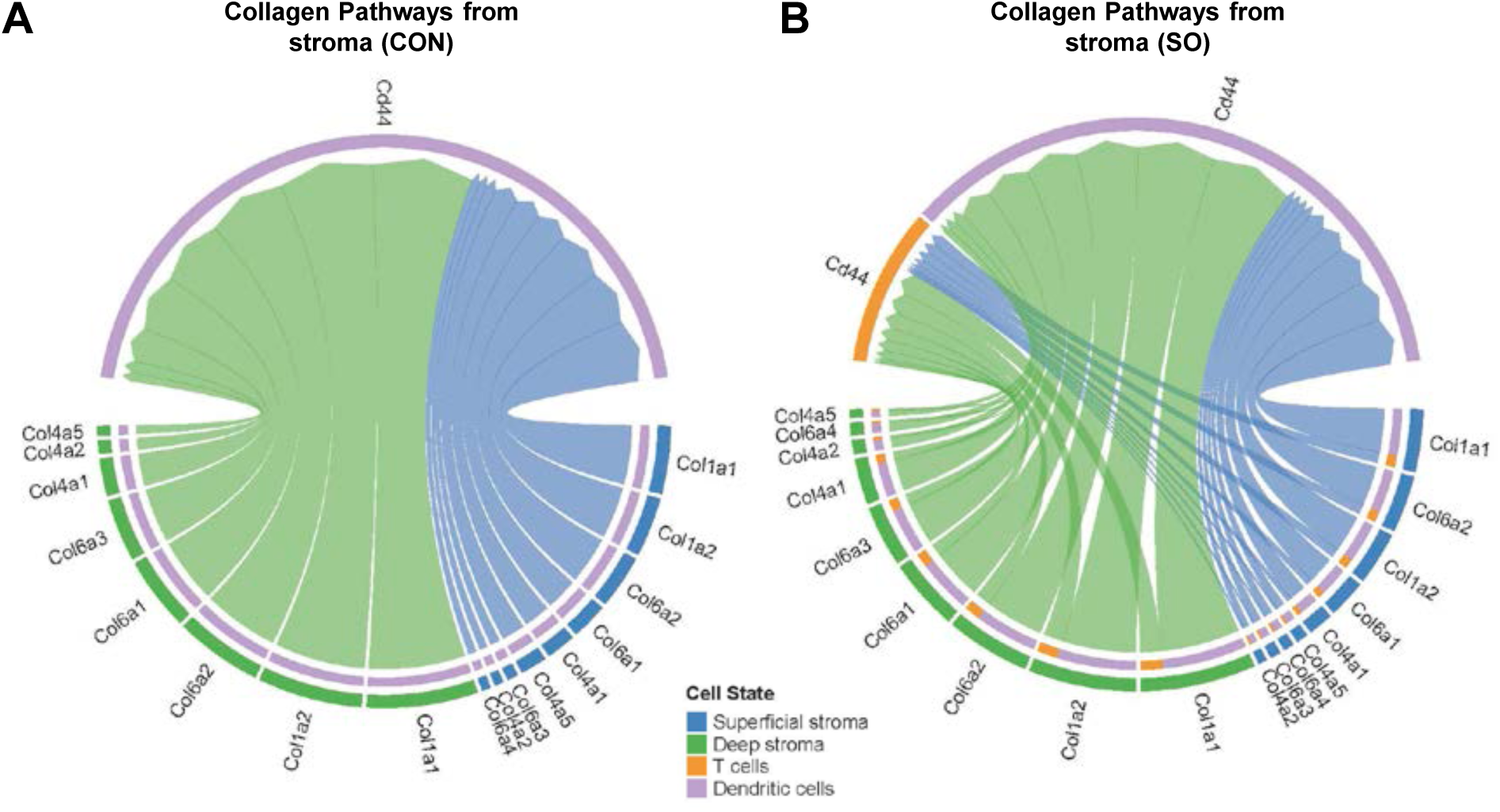
Collagen ligand-receptor interactions between stromal and immune compartments are expanded in the superovulated uterus. **(A-B)** Chord diagrams depicting collagen ligand-receptor (*Col–Cd44*) signaling received by dendritic cells and T cells from the superficial and deep stroma in control **(A)** and superovulated **(B)** uteri. In control, collagen signaling is predominantly received by dendritic cells from both stromal sub-clusters. During superovulation, collagen signaling additionally engages T cells, with increased interaction involving *Col1a1*, *Col1a2*, *Col4a1*, *Col4a2*, *Col4a5*, *Col6a1*, *Col6a2*, *Col6a3*, and *Col6a4*. Arc size represents the relative signaling strength of each ligand-receptor interaction. CON, control; SO, superovulation.

**Supplementary Figure 9.**
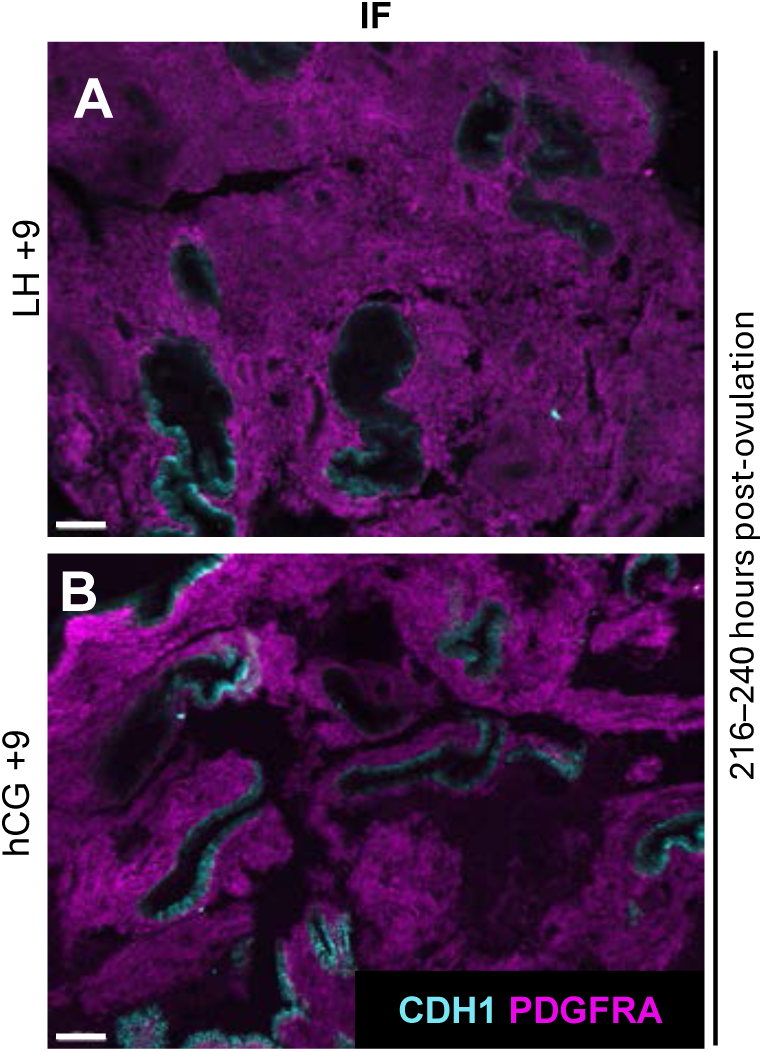
Comparable PDGFRA expression observed in the stimulated and normal cycling human uterus during the window of implantation. **(A-B)** PDGFRA immunofluorescence staining of human uterus biopsy samples from normal cycling women, LH+9 **(A)**, and stimulated women, hCG+9 **(B)**, around the window of implantation. LH+9, control (human); hCG+9, superovulation (human). Scale bars: **A,B:** 100 µm.

**Supplementary Figure 10.**
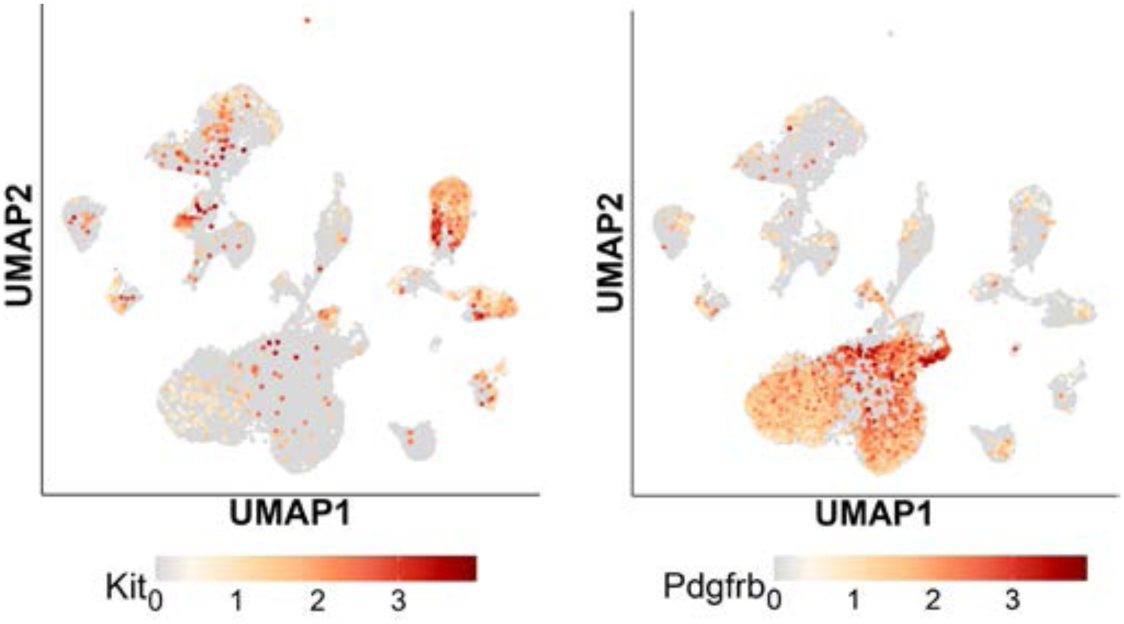
*Kit* expression is minimal in the stromal and smooth muscle compartments, and *Pdgfrb* is expressed in the stroma at GD3 0900 h in the scRNA-seq dataset.

**Supplementary Figure 11.**
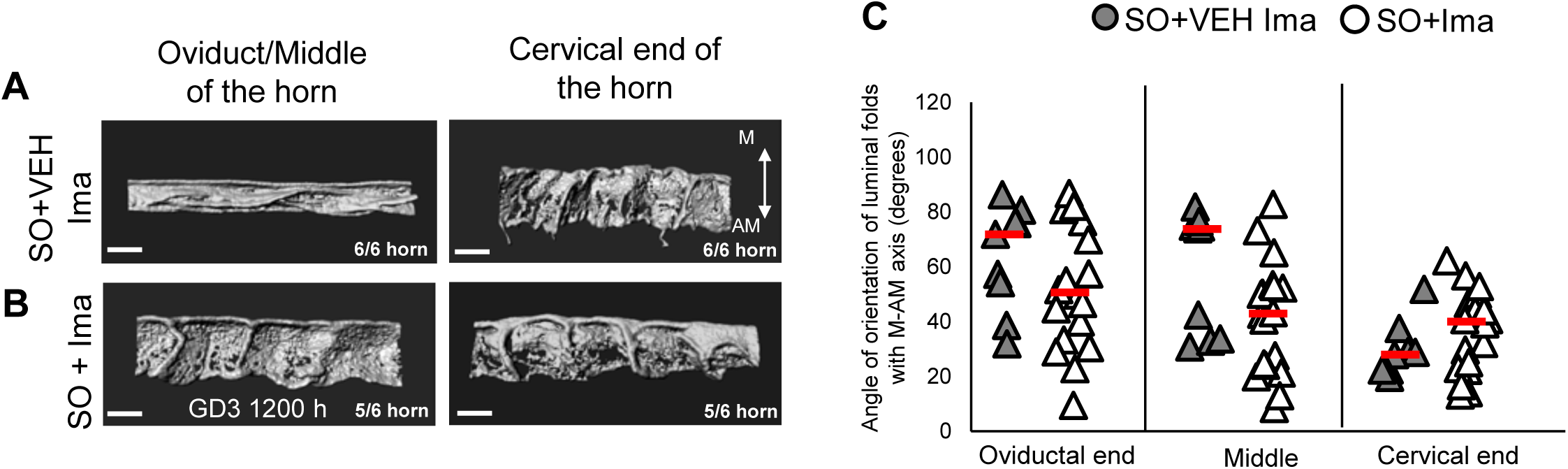
Longitudinal folds in the oviductal and middle regions during superovulation are rescued with Imatinib treatment. **(A-B)** 3D reconstructions of the uterine lumen in superovulation+vehicle **(A)** and superovulation+imatinib **(B)** uteri at GD3 1200 h. Longitudinal folds are present in the oviductal and middle regions of the uterine horn in superovulation+vehicle; however, superovulation+imatinib displays transverse folding throughout the uterine horn similar to control. **(C)** Quantification of fold angle relative to the M–AM axis, comparing the oviductal, middle and cervical regions of the uteri at GD3 1200 h (n = 3 mice/group). CON, control; SO+VEH Ima, vehicle for imatinib; SO+Ima, superovulated mice treated with imatinib. Scale bars: **A-B:** 300 µm.

**Supplementary Figure 12.**
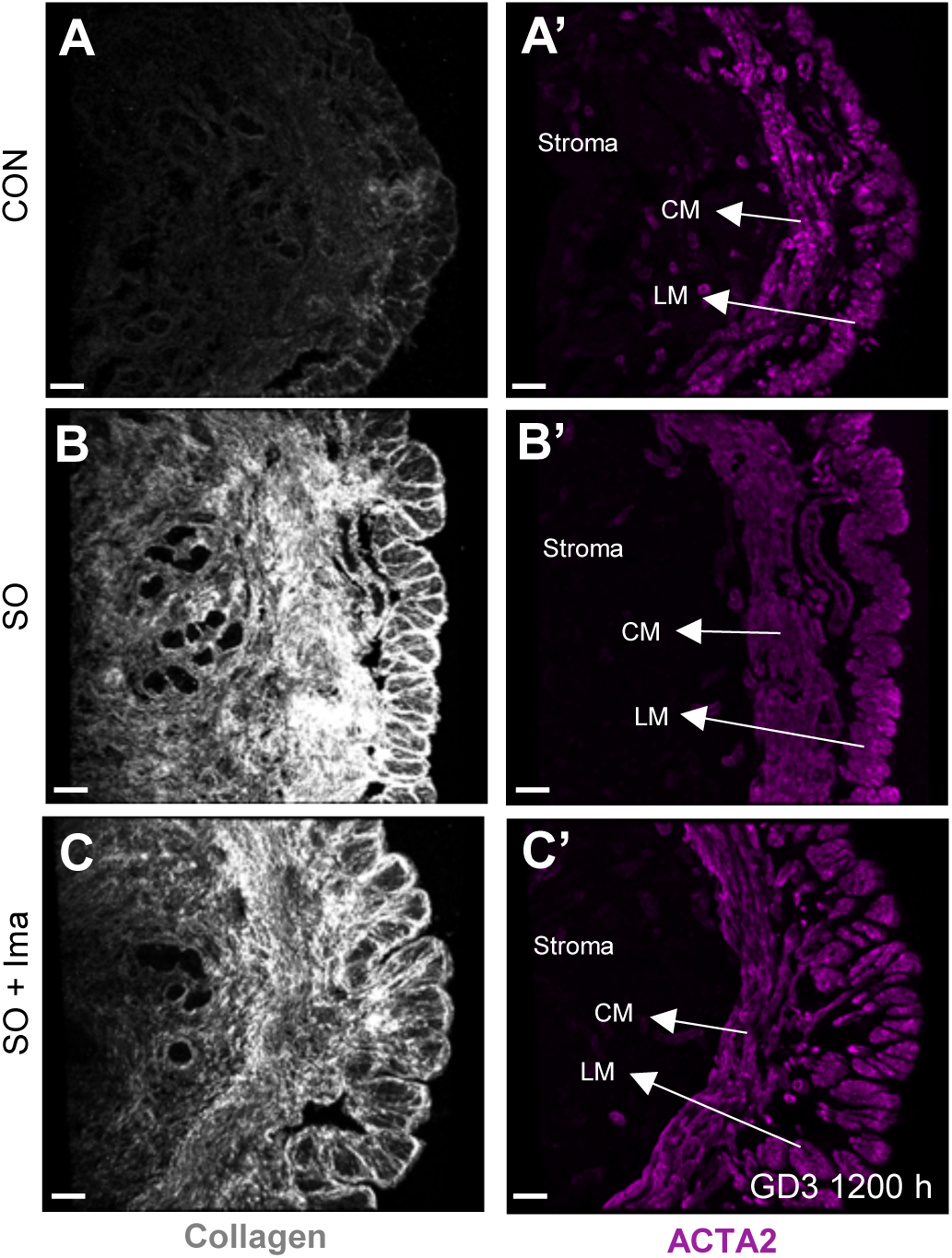
Short-term imatinib treatment does not rescue excess collagen deposition in the superovulated uterus. **(A-C)** Second harmonic generation imaging of fibrillar collagen in the stroma and smooth muscle of control **(A)**, superovulation **(B)**, and superovulation+imatinib **(C)** at GD3 1200 h. Enhanced collagen deposition is observed in superovulation **(B)** and superovulation+imatinib (n=3 mice/group; n_t_ = 9 transverse sections/mice) **(C)** uteri compared to control. **(A’-C’)** ACTA2 immunofluorescence staining corresponding to **A**, **B**, and **C**. n_t_, number of transverse sections; CON, control; SO, superovulation; SO+Ima, superovulated mice treated with imatinib; CM, circular smooth muscle; LM, longitudinal smooth muscle. Scale bars: **A-C,A’-C’:** 50 µm.

**Supplementary Figure 13.**
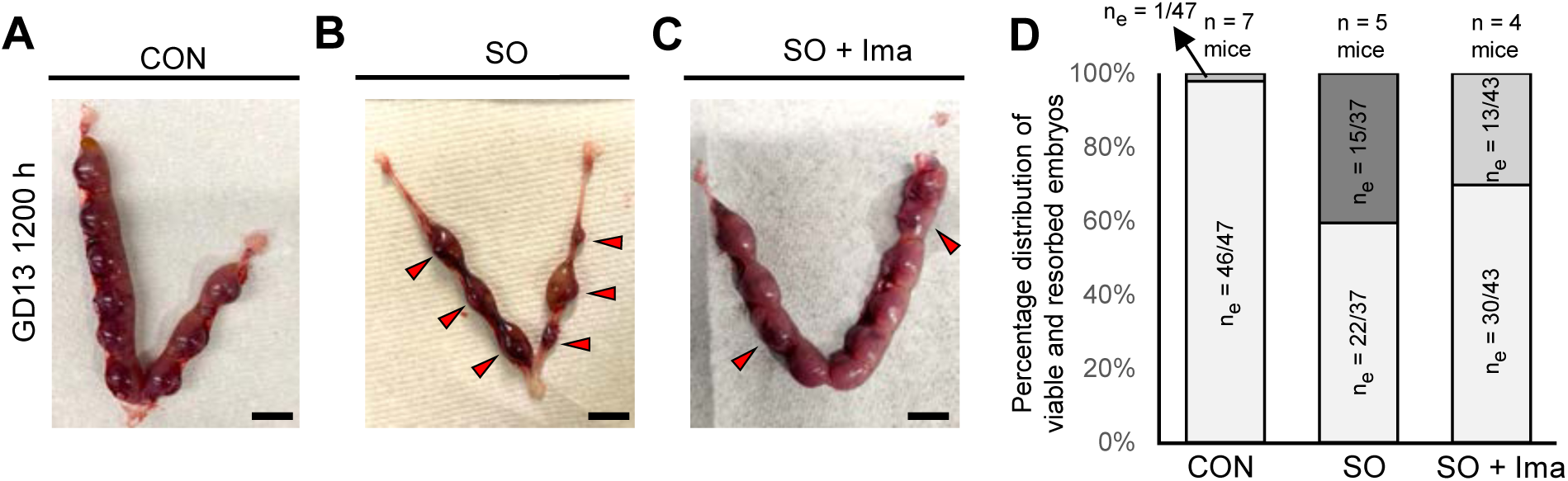
Mid-gestation embryonic development in superovulated mice treated with PDGFRA inhibitor. **(A-C)** Uterine horns with embryos at GD13 1200 h in control **(A)**, superovulation **(B)** and superovulation+imatinib **(C)** uteri (control and superovulation: n = 5 mice/group; superovulation+imatinib: n = 4 mice). Red arrowheads: resorption sites. Embryo viability is assessed by the presence of a fetal heartbeat. **(D)** Percentage distribution of viable and resorbed embryos in control (n = 7 mice, n_e_ = 47 embryos), superovulation (n = 5 mice, n_e_ = 37 embryos), and superovulation+imatinib (n = 4 mice, n_e_ = 43 embryos) uteri at GD13 1200 h. Control uteri display a high proportion of viable embryos (97.87%); superovulated uteri show a reduction in viability (59.45%); superovulation+imatinib uteri show a partial recovery in viability (69.76%). n_e_, number of embryos; CON, control; SO, superovulation; SO+Ima, superovulated mice treated with imatinib. Scale bars: **A-C:** 5mm.

**Supplementary Table 1.** Superovulation results in a progressive decrease in viable embryos between GD5 1200 h and GD13 1200 h. At GD4 1800h, 54.68% of embryos in superovulated mice display a V-shaped chamber with proper alignment of the embryo-uterine axis, whereas 39.06% are trapped in a fold (Chtrap) and 6.25% have no chamber (Chnull). At GD5 1200h, embryos display abnormal implantation chambers that do not resemble the elongated V-shaped morphology seen in controls, including no chamber (Chnull), short chamber (Chshort), and tilted chamber (Chtilt), as described in Fig. 2K; notably, embryos in the Chshort category appear morphologically normal. At GD13 1200h, 40.54% of embryos are resorbed. Accounting for both deformed and resorbed embryos, the total proportion of viable embryos drops to approximately 34.37% in superovulated mice at mid-gestation. CON, control; SO, superovulation.

| Gestational day | CON |  |  |  |  | SO |  |  |  |  |
| --- | --- | --- | --- | --- | --- | --- | --- | --- | --- | --- |
|  | Normal (n) | Abnormal (ab) | Total (n<sub>tot</sub>) | Proportion of embryo loss (ab/n<sub>tot</sub>)*100 | Percentage of viable embryos based on n<sub>tot</sub> at GD4 1800h (n/n<sub>tot</sub> GD4 1800h)*100 | Normal (n) | Abnormal (ab) | Total (n<sub>tot</sub>) | Proportion of embryo loss (ab/n<sub>tot</sub>)*100 | Percentage of viable embryos based on n<sub>tot</sub> at GD4 1800h (n/n<sub>tot</sub> GD4 1800h)*100 |
| GD4 1800h | 32 | 0 | 32 (n=4)<br>Avg - 8 | 0% | 100% | 35 | (Ch<sub>trap</sub>) In fold – 25<br>(Ch<sub>null</sub>) No chamber – 4 | 64 (n=4)<br>Avg-16 | ~45.3% | ~54.68% |
| GD5 1200h | 31 | 0 | 31 (n=4)<br>Avg- 7.75 | 0% | ~100% | (Ch<sub>norm</sub>) Elongated chamber - 12<br>(Ch<sub>short</sub>) Short chamber – 18 | (Ch<sub>null</sub>) No implantation chamber - 6<br>(Ch<sub>tilt</sub>) tilted chamber - 21 | 57 (n=4)<br>Avg- 14.25 | ~47.4% | ~46.87% |
| GD13 1200h | 42 | 1 | 43 (n=5)<br>Avg- 8.6 | 2.33% | ~100% | 22 | Resorbed - 15 | 37 (n=5)<br>Avg- 7.4 | ~40.54% | ~34.37% |

**Supplementary Table 2.** Overall volume of PDGFRA-expressing cells normalized to Hoechst at GD 1 1200h and GD3 1200h. Mann–Whitney U-test *p*-values for the overall volume of cells expressing PDGFRA and Hoechst in stromal and smooth muscle compartments at GD1 1200h and GD3 1200h, comparing SO and CON mice. No significant differences are detected in either compartment at GD1 1200 h or GD3 1200 h. CON, control; SO, superovulation.

|  | GD1 1200h |  |  |  | GD3 1200h |  |  |  |
| --- | --- | --- | --- | --- | --- | --- | --- | --- |
|  | Stroma |  | Smooth muscle |  | Stroma |  | Smooth muscle |  |
|  | Median PDGFRA+ volume / Hoechst volume | *p*-value | Median PDGFRA+ volume / Hoechst volume | *p*-value | Median PDGFRA+ volume / Hoechst volume | *p*-value | Median PDGFRA+ volume / Hoechst volume | *p*-value |
| CON | 0.9684 | 0.2063 | 0.8049 | 0.5360 | 1.0962 | 0.6837 | 0.8873 | 0.9351 |
| SO | 1.0061 |  | 0.8474 |  | 1.1518 |  | 0.9255 |  |

**Supplementary Table 3.** Progesterone levels in humans and mice. Data are shown as mean ± SD. Mann-Whitney U test. Human: LH+2, control; hCG+2, superovulation; Mice: CON, control; SO, superovulation.

|  | Human biopsy samples (peri-ovulatory (Chemerinski et al., 2024)) |  |  | Mouse pre-implantation (GD3 1200h) |  |  |
| --- | --- | --- | --- | --- | --- | --- |
|  | LH +2 (n=10) | hCG +2 (n=13) | *p*-value | CON (n=5) | SO (n=8) | *p*-value |
| P4 (ng/mL) | 1.0±0.7 | 12.1±5.3 | <0.0001 | 24.7±8.6 | 81.7±32.5 | <0.001 |

